# Low intensity repetitive transcranial magnetic stimulation enhances remyelination by newborn and surviving oligodendrocytes in the cuprizone model of toxic demyelination

**DOI:** 10.1101/2024.02.29.582855

**Authors:** Phuong Tram Nguyen, Kalina Makowiecki, Thomas S Lewis, Alastair Fortune, Mackenzie Clutterbuck, Laura Reale, Bruce V Taylor, Jennifer Rodger, Carlie L Cullen, Kaylene M Young

**Affiliations:** Menzies Institute for Medical Research, University of Tasmania, Hobart, TAS Australia; School of Biological Sciences, The University of Western Australia, Crawley, WA, Australia; Perron Institute for Neurological and Translational Science, Nedlands, WA, Australia; Mater Research Institute, The University of Queensland, Woolloongabba, QLD, Australia

**Keywords:** demyelination, remyelination, surviving oligodendrocyte, oligodendrocyte, internode, cortex, corpus callosum, repetitive transcranial magnetic stimulation (rTMS), OLIG2, ASPA

## Abstract

In people with multiple sclerosis (MS), newborn and surviving oligodendrocytes (OLs) can contribute to remyelination, however, current therapies are unable to enhance or sustain endogenous repair. Low intensity repetitive transcranial magnetic stimulation (LI-rTMS), delivered as an intermittent theta burst stimulation (iTBS), increases the survival and maturation of newborn OLs in the healthy adult mouse cortex, but it is unclear whether LI-rTMS can promote remyelination. To examine this possibility, we fluorescently labelled oligodendrocyte progenitor cells (OPCs; *Pdgfr*α*-CreER* transgenic mice) or mature OLs (*Plp-CreER* transgenic mice) in the adult mouse brain and traced the fate of each cell population over time. Multiple consecutive daily sessions of iTBS (600 pulses; 120 mT), delivered during cuprizone (CPZ) feeding, did not alter new or pre-existing OL survival but increased the number of myelin internodes elaborated by new OLs in the primary motor cortex (M1). This resulted in each new M1 OL producing ∼471µm more myelin. When LI-rTMS was delivered after CPZ withdrawal (during remyelination), it significantly increased the length of the internodes elaborated by new M1 and callosal OLs and increased the number of surviving OLs that contributed to remyelination in the corpus callosum (CC). As LI-rTMS can non-invasively promote remyelination by modifying the behaviour of new and surviving OLs, it may be suitable as an adjunct intervention to enhance remyelination in people with MS.

## Introduction

Oligodendrocytes (OLs) elaborate and wrap myelin around axons in the central nervous system (CNS), effectively increasing the resistance and decreasing the capacitance of the axonal membrane, to enable the saltatory conduction of action potentials [1]. OLs also detect neuronal activity and provide metabolic support to the axons they myelinate [2]. OL loss and demyelination contribute to neurodegeneration and disability accrual in people with multiple sclerosis (MS) [3]. In mice and humans, oligodendrogenesis and spontaneous remyelination can occur in response to a demyelinating injury [4–7]. In people with MS, the level of endogenous remyelination is highly variable and its eventual failure has been largely attributed to the inability of OPCs to differentiate into new OLs within the lesion environment [8, 9] and surviving OLs to generate new myelin sheaths [7, 10]. Therapeutic approaches designed to enhance remyelination are most likely to be effective if they simultaneously support repair by new and surviving OLs.

Neuronal activity is a major extrinsic regulator of myelination [11–13]. Repetitive transcranial magnetic stimulation (rTMS) applies focal magnetic fields to induce electric currents in the brain and can be used to noninvasively modulate neuronal activity, with the outcome being influenced by the stimulation intensity, frequency, pattern and number of sessions [14, 15]. rTMS has been delivered to people with MS to evaluate its effect on MS clinical signs, including impaired hand dexterity [16, 17], lower urinary tract function [18, 19] and working memory [20], spasticity [21–23] or fatigue [24]. However, these clinical studies did not explore the impact of rTMS on remyelination. We previously reported that 28 consecutive daily sessions of low-intensity (LI)-rTMS, patterned as an intermittent theta burst stimulation (iTBS), increased the survival and maturation of new OLs. LI-rTMS not only increased the number of new OLs added to the adult mouse cortex but increased the length of the internodes the new OLs produced [25]. There is also evidence that rTMS can protect against demyelination or enhance remyelination, as 14 consecutive daily sessions of 10 Hz rTMS (0.4 T) can reduce the level of gross demyelination detected following a spinal cord injury [26], and 7 consecutive daily sessions of 5Hz rTMS (1.26 T) can reduce the level of hippocampal, cortical and striatal demyelination detected after cuprizone (CPZ)-feeding [27].

To determine whether LI-rTMS can influence the behaviour of new and / or surviving OLs in the demyelinating or remyelinating CNS, we performed cre-lox lineage tracing to follow the fate of OPCs and the new OLs they generate (*Pdgfr*α *-CreER* transgenic mice), or the mature myelinating OLs that survive demyelination (*Plp-CreER* transgenic mice) in the adult mouse brain. When LI-rTMS was delivered as an iTBS during CPZ feeding, new M1 OLs elaborated significantly more internodes, than OLs in sham-stimulated mice. Furthermore, when iTBS was delivered after CPZ withdrawal, it increased the length of internodes generated by new M1 OLs and increased the proportion of surviving callosal OLs that contributed to remyelination.

## Materials and methods

### Transgenic mouse genotyping and tamoxifen delivery

All animal experiments were approved by the University of Tasmania Animal Ethics (A0018606) and carried out in accordance with the Australian code of practice for the care and use of animals for scientific purposes. *Pdgfr*α*-CreERT^TM^* (RRID: IMSR_JAX:018280), *Plp-CreER^T^* (RRID: IMSR_JAX:005975), *Rosa26-YFP* (RRID: IMSR_JAX:006148), and *Tau-mGFP* (RRID: IMSR_JAX:021162) mouse lines were purchased from the Jackson Laboratories. *Pdgfr*α*-CreERT^T2^* transgenic mice [28] were a kind gift from Prof. William D Richardson (University College London). All transgenic mice were maintained on a C56BL/6J background.

To fluorescently label and trace OPCs and the new OLs they produce, heterozygous *Pdgfr*α*-CreERT^TM^* transgenic mice [29] were crossed with homozygous *Rosa26-YFP* Cre-sensitive reporter mice [30] or heterozygous *Pdgfr*α*-CreERT^T2^* transgenic mice [28] were crossed with heterozygous *Tau-lox-STOP-lox-mGFP-IRES-NLS-LacZ-pA* (*Tau-mGFP*) Cre-sensitive reporter mice [31] to generate double heterozygous offspring. To fluorescently label and trace pre-existing mature OLs, heterozygous *Plp-CreER^T^* transgenic mice [32] were crossed with homozygous *Rosa26-YFP* or heterozygous *Tau-mGFP* Cre-sensitive reporter mice to generate double heterozygous offspring. The expression of *Cre*, *Rosa26-YFP* or *mGFP* transgenes was confirmed by polymerase chain reaction (PCR) as described [25]. In brief, genomic DNA (gDNA) was extracted from ear biopsies by ethanol precipitation and PCR performed using 50–100 ng of gDNA with the following primer combinations: Cre 5′ CAGGT CTCAG GAGCT ATGTC CAATT TACTG ACCGTA and Cre 3′ GGTGT TATAAG CAATCC CCAGAA; Rosa26 wildtype 5′ AAAGT CGCTC TGAGT TGTTAT, Rosa26 wildtype 3′ GGAGC GGGAG AAATG GATATG and Rosa26 YFP 5′ GCGAA GAGTT TGTCC TCAACC; GFP 5′ CCCTG AAGTTC ATCTG CACCAC and GFP 3′ TTCTC GTTGG GGTCT TTGCTC. Alternatively, mice were genotyped by quantitative PCR (Transnetyx).

Experimental mice were group-housed with same-sex littermates (2-5 per cage) in Optimice microisolator cages on a 12 h light/dark cycle (lights on 7:00, lights off 19:00) at 21 ± 2°C with *ad libitum* access to standard rodent chow (Barrastoc rat and mouse pellets) and water. Mice weighed 16-24 g at the start of the experiment, when male and female mice were randomly assigned to each treatment. Care was taken to ensure littermates and sexes were represented across treatment groups. To initiate Cre-mediated recombination of a reporter transgene, tamoxifen (Tx; Sigma, T5648) was dissolved in corn oil (Sigma, C8267) at a concentration of 40 mg/mL by sonication at 21°C for 2 hours and administered to P60 mice (P60 ± 5 days) by oral gavage at a dose of 300 mg/kg body weight daily for 4 consecutive days.

### CPZ-induced demyelination and remyelination

A diet containing 0.2% (w/w) CPZ (Sigma, C9012) was fed to mice for 4 or 5 weeks, as specified, from P67 (P67± 5 days). CPZ was thoroughly mixed into ground rodent chow and 1mL of H_2_O added per 2g dry weight. Approximately 25 g of hydrated CPZ chow was prepared per mouse, replaced every second day. Mice were transferred to clean home cages every 4 days to prevent them from ingesting old CPZ chow.

### LI-rTMS

LI-rTMS was delivered as previously described [25] using a custom made 120mT circular coil designed for rodent stimulation (8 mm outer diameter, iron core) [33]. iTBS consisted of bursts of 3 pulses at 50 Hz, repeated at 5 Hz for a 2-s period (10 bursts), followed by an 8-s gap, repeated for 20 cycles (total 600 pulses, 192 s). Stimulation parameters were controlled by a waveform generator using custom monophasic waveforms (400µs rise time; Agilent Benchlink Waveform Builder), connected to a bipolar voltage programmable power supply, at maximum power output (100V) (KEPCO BOP 100-4M, TMG test equipment). Mice were habituated to the stimulation room for 1 h prior to delivering a sham stimulation or iTBS, and the intervention was delivered once per day, at the same time, for the specified duration (2 or 4 weeks). For the ∼3 min required, mice were gently restrained using plastic body-contour shape restraint cones (0.5 mm thick; Able Scientific). The coil was manually held over the midline of the head with the back of the coil positioned in line with the front of the ears (∼ Bregma –3.0). For sham stimulation, the procedure was the same, but no current was passed through the coil.

### Tissue preparation and immunohistochemistry

Mice were terminally anesthetised with an intraperitoneal injection of sodium pentobarbitone (150 mg/kg body weight) and were transcardially perfused with 4% (w/v) paraformaldehyde (PFA; Sigma) in phosphate buffered saline (PBS; pH 7.4). Brains were cut into 2 mm-thick coronal slices using a brain matrix (Kent Scientific) before being post-fixed in 4% PFA in PBS at 21°C for 1.5-2h. Tissue was cryoprotected overnight in 20% (w/v) sucrose (Sigma) in PBS at 4°C, then embedded in OCT (optimal cutting temperature matrix; Thermo Fisher Scientific) and snap frozen in liquid nitrogen and stored at -80°C.

Coronal brain cryosections (30 µm) containing M1 (∼ Bregma +0.5) or V2 (∼ Bregma –2.5) were collected and processed as floating sections [34]. Sections were pre-incubated in blocking solution [10% (v/v) fetal calf serum, 0.1% Triton X-100 in PBS, pH 7.4] for 1h at 21°C on an orbital shaker. Primary and secondary antibodies were diluted in blocking solution and applied to cryosections overnight at 4°C on an orbital shaker, unless staining involved the use of mouse anti-CASPR (Clone K65/35; 1:200, NeuroMab 75-001), in which case sections were incubated in primary antibodies for 48 hours at 4°C. Primary antibodies included goat anti- PDGFRα (1:100, R&D Systems AF1062), rabbit anti-OLIG2 (1:400, Millipore AB9610), rabbit anti-ASPA (1:200, Abcam ab97454) and rat anti-GFP (1:2000, Nacalai Tesque 04404-26). Secondary antibodies, conjugated to AlexaFluor-488, -568 or -647 (Invitrogen) included donkey anti-goat (1:1000), donkey anti-rabbit (1:1000), donkey anti-mouse (1:1000) and donkey anti-rat (1:500). Nuclei were labelled using Hoechst 33342 (1:1000, Invitrogen). After washing in PBS, floating sections were mounted onto glass slides (Superfrost) and coverslipped with fluorescent mounting medium (Dakomount, Agilent Technologies).

### Black gold II myelin staining

Black gold II myelin staining was performed using the Biosensis Black gold II RTD kit (Sapphire Biosciences, TR-100-BG) according to the manufacturer’s instructions. Briefly, 30 µm coronal brain cryosections from ∼ Bregma +0.5 were mounted on slides and allowed to dry before being hydrated in MilliQ water for 2 min. Slides were incubated with preheated black gold II (diluted 1:10 in MilliQ water) in the dark at 65°C for 25 min, then rinsed in MilliQ water for 2 min before incubating with sodium thiosulfate (diluted 1:10) for 3 min. Slides were washed in MilliQ water (3 x 5 min), and dehydrated using a series of graded alcohol steps, before clearing with xylene (Sigma, 214736) for at least 2 min, and mounted with DPX mounting medium (Sigma, 06522).

### Microscopy and cell quantification

Confocal images were obtained using an UltraView Nikon Ti Microscope with Volocity software (Perkin Elmer, Massachusetts, United States). For quantification of cell number, low magnification (20 × objective) images were taken spanning M1, V2, or the corpus callosum (CC), underlying M1. The z stack images (3 µm z-spacing) were collected using standard excitation and emission filters for DAPI, FITC (AlexaFluor-488), TRITC (AlexaFluor-568), and CY5 (AlexaFluor-647) and stitched together to make a composite image of the defined region of interest. To quantify internode number per mGFP^+^ OL and to measure the length of individual mGFP^+^ internodes, each mGFP^+^ OL with a cell body situated in M1 or V2 was positioned in the centre of the field of view and a high magnification image (40 × objective) collected with 0.5 µm z-spacing. To quantify mGFP^+^ internode length in the CC, we defined a region encompassing an mGFP^+^ cell body and surrounding internodes that was imaged at a high magnification (60 × objective) with 0.5 µm z-spacing and stitched together to form a single image for analysis. To quantify black gold II myelin labelling, we collected low magnification (2.5x objective) images, using a light microscope with Zeiss software, that included M1 and the underlying CC.

Cell counts or internode number and length measurements were performed manually in ImageJ (NIH) or Photoshop CS6 (Adobe, San Jose, United States). The researcher performing the measurements was blind to the experimental group and treatment conditions. The accuracy of quantification was verified across a subset of images by a second researcher, who was also blind to the experimental group and treatment conditions. Cell counts were carried out according to predetermined criteria. YFP^+^ or mGFP^+^ cells were only quantified when they contained a Hoechst 33342^+^ nucleus. YFP^+^ or mGFP^+^ cells were classified as PDGFRα^+^ OPCs or ASPA^+^ OLs, when the PDGFRα or ASPA fluorescent labelling was within the bounds of the YFP^+^ / mGFP^+^ cell borders. YFP^+^ or mGFP^+^ cells were classified as OLIG2^+^ when OLIG2 labelling overlapped with the Hoechst 33342^+^ nucleus. The length of an individual M1, V2 or CC mGFP^+^ internode was only measured when the internode was flanked by CASPR^+^ paranodes. Within M1 and V2, the low density of mGFP^+^ OLs allowed internodes to be discerned, counted, or measured, and attributed to individual OLs. The high density of mGFP^+^ OLs in the CC prevented individual mGFP^+^ internodes from being assigned to a specific mGFP^+^ OL. Therefore, the length of individual CASPR-flanked mGFP^+^ internodes was measured, but the number of internodes per mGFP^+^ CC OL was not quantified.

The number of mice in each group or the number of OLs or internodes analysed (n) is stated in each figure legend. When quantifying cell density from maximum projection images, the total number of cells within the defined region was divided by the x-y area and expressed as cells per mm^2^ (not corrected for z-depth, but tissue sections were 30µm). We calculated the proportion (%) of YFP^+^ or mGFP^+^ cells that were OLs using the following formulae: (the number of YFP^+^ PDGFRα-neg OLIG2^+^ OLs within the defined region of interest) / (the total number of YFP^+^ cells in the region of interest) x 100. To quantify the proportion (%) of mGFP^+^ cells that were PDGFRα^+^ OLIG2^+^ OPCs or had matured into PDGFRα-neg OLIG2^+^ premyelinating or myelinating OLs, we applied the following formulae: (the number of mGFP^+^ cells of each subtype in the region of interest) / (the number of mGFP^+^ cells in the region of interest) x 100. To quantify the proportion (%) of mGFP^+^ surviving OLs with or without myelin, that were OLIG2^+^ or OLIG2-neg, the number of cells in each category was divided by the total number of mGFP^+^ cells x 100. To approximate the length of myelin sheath synthesized by individual mGFP^+^ OLs (µm), average internode length for the mGFP^+^ OL was multiplied by the number of internodes elaborated by that mGFP^+^ OL.

### Statistical analyses

All statistical analyses were performed using Prism 8 (GraphPad Software). When data were normally distributed, they were analysed using an unpaired two-tailed t-test, and when data were not normally distributed, they were analysed using a Mann-Whitney U test, as specified in the figure legends. When internode length, internode number or predicted total myelin data were analyzed per OL, data were nested within mice and analysed using a nested t-test. Internode length cumulative distributions were analysed using a Kolmogorov-Smirnov (KS) test. When comparing cell counts across treatment conditions, but also between anatomical regions of the same mice, data were analysed using mixed two-way ANOVAs, with region as the repeated measures factor (Greenhouse-Geisser corrected if sphericity was violated), and treatment condition as the between-subjects factor. Follow-up tests for simple effects of treatment, within each brain region, were Bonferroni corrected for multiple comparisons. To determine whether the mean differs from 0, data for each group were analysed separately using a one sample t-test or Wilcoxon signed-rank test, to compare the group mean or median, respectively, to 0. Multiple comparisons were corrected (with a false discovery rate set at 5%) using the Benjamini-Hochberg method. Statistical significance was defined as p < 0.05, corrected for multiple testing. Further statistical information is provided in each figure legend including, for example, ANOVA main effects.

### Data availability

The image files and data described in this manuscript can be obtained from the corresponding author by reasonable request.

## Results

### iTBS delivered during CPZ feeding does not alter the number of new OLs added to the cortex or CC

In the CPZ model of demyelination, OPCs proliferate and generate new OLs that rapidly differentiate upon CPZ withdrawal [6]. To determine whether iTBS, delivered during CPZ feeding and in the first week following CPZ withdrawal, can increase the number of new OLs added to the cortex or CC, Tx was administered to P60 *Pdgfr*α*-CreERT^TM^ :: Rosa26-YFP* transgenic mice to fluorescently label OPCs throughout the CNS (**Fig. 1a**). From P67, *Pdgfr*α*-CreERT^TM^ :: Rosa26-YFP* mice were either maintained on a control diet as treatment naïve mice (P67+42) or transferred to a CPZ diet for 35 days to induce demyelination and returned to control chow for a further 7 days to enable remyelination (P67+35CPZ+7). On day 14 of CPZ feeding, mice commenced sham stimulation or iTBS, which was delivered daily for 28 consecutive days i.e., the remainder of the 42-day time-course (**Fig. 1a**). Coronal brain cryosections were immunolabelled to detect YFP, PDGFRα (OPCs) and OLIG2 (all cells of the OL lineage) (**Fig. 1b-g**) in M1, V2 and the CC. These regions were selected for analysis as they are demyelinated by CPZ-feeding [6, 35–37] and OLs within these regions respond to LI-rTMS in healthy mice [25, 38]. We found that >95% of PDGFRα^+^ parenchymal M1 and V2 OPCs were YFP-labelled in P67+42 naïve and P67+35CPZ+7 sham or iTBS treated *Pdgfr*α*-CreERT^TM^ :: Rosa26-YFP* mice (**Fig. 1h**). Total M1 or V2 OPC density was also equivalent in P67+42 naïve and P67+35CPZ+7 sham or iTBS treated mice (**Fig. 1i**). While >95% of PDGFRα^+^ OPCs were similarly YFP-labelled in the CC of P67+42 naïve *Pdgfr*α*-CreERT^TM^ :: Rosa26-YFP* mice, reflecting the initial recombination efficiency (**Fig. 1h**). Fewer OPCs were YFP-labelled in the CC after CPZ-feeding, with only ∼66% of OPCs being YFP-labelled in the CC of P67+35CPZ+7 iTBS mice (**Fig. 1h, j, k**), presumably reflecting the generation of YFP-neg PDGFRα^+^ OPCs by subventricular zone neural stem cells in response to callosal demyelination [39, 40]. As the proportion of callosal OPCs that is YFP- labelled is equivalent in P67+35CPZ+7 sham and iTBS *Pdgfr*α*-CreERT^TM^ :: Rosa26-YFP* mice (**Fig. 1h**), iTBS does not enhance OPC generation from neural stem cells. Furthermore, as total PDGFRα^+^ callosal OPC density was equivalent in naïve and P67+35CPZ+7 sham or iTBS mice (**Fig. 1i**), iTBS does not affect OPC density in this region.

**Figure 1:**
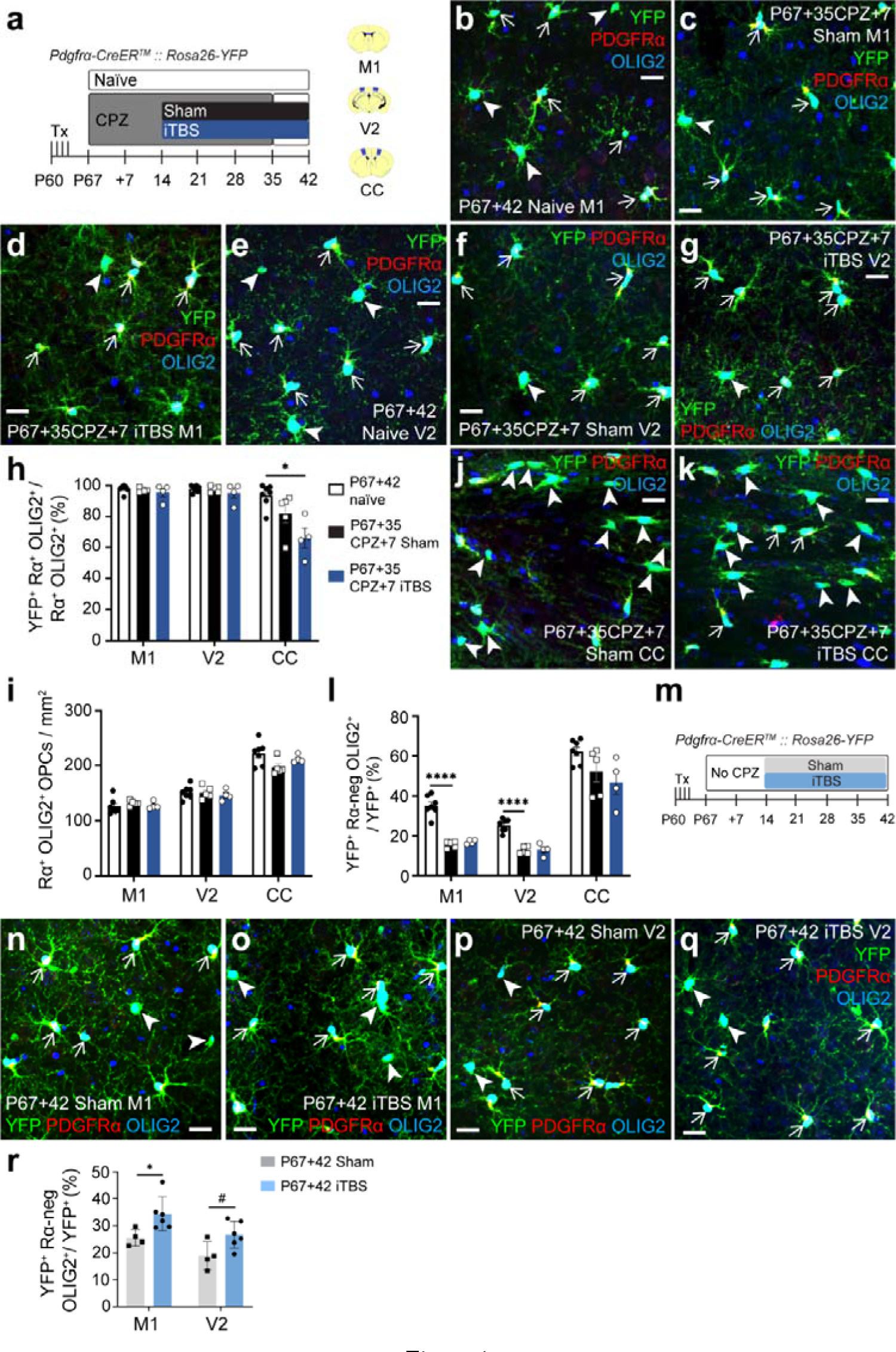
*iTBS during demyelination does not increase new OLs added to the cortex and corpus callosum.* **a.** Experimental schematic showing the timeline over which *Pdgfr*α*-CreERT^TM^ :: Rosa26-YFP* mice received 4 consecutive days of Tx gavage, up to 35 days of CPZ and up to 28 days of LI-rTMS (sham-stimulation or iTBS). This experiment included three treatment groups: P67+42 naïve, P67+35CPZ+7 sham, and P67+35CPZ+7 iTBS. The brain regions analysed are shown in blue in the coronal brain section schematics: M1 (∼ Bregma +0.5), V2 (∼ Bregma -2.5) and the CC (∼ Bregma +0.5). **b-g.** Confocal images of YFP (green), PDGFRα (red) and OLIG2 (blue) immunohistochemistry in M1 (**b-d**) and V2 (**e-g**) of P67+42 naïve (**b, e**), P67+35CPZ+7 sham (**c, f**) and P67+35CPZ+7 iTBS (**d, g**) mice. YFP^+^ PDGFRα^+^ OPCs are denoted by arrows. YFP^+^ PDGFRα-neg OLs are denoted by arrowheads. **h.** The proportion (%) of PDGFRα^+^ OLIG2^+^ OPCs that expressed YFP in the M1, V2 and CC of P67+42 naïve (white, n=7), P67+35CPZ+7 sham (black, n=5) or P67+35CPZ+7 iTBS (blue, n=4) mice. Repeated measures two-way ANOVA with Geisser-Greenhouse correction: treatment F (2, 13) = 5.294, p = 0.021; region F (1.027, 13.35) = 48.37, p < 0.0001; interaction F (4, 26) = 9.709, p < 0.0001. **i.** Density of PDGFRα^+^ OLIG2^+^ OPCs in M1, V2 and the CC of P67+42 naïve (white, n=7), P67+35CPZ+7 sham (black, n=5) or P67+35CPZ+7 iTBS (blue, n=4) mice. Repeated measures two-way ANOVA with Geisser-Greenhouse correction: treatment F (2, 13) = 0.9164, p = 0.4243; region F (1.943, 25.26) = 188.5, p < 0.0001; interaction F (4, 26) = 2.822, p = 0.0454. **j-k.** Confocal images of YFP (green), PDGFRα (red) and OLIG2 (blue) immunohistochemistry in the CC of P67+35CPZ+7 sham (**j**) and P67+35CPZ+7 iTBS (**k**) mice. **l.** The proportion (%) of YFP^+^ cells that are newly differentiated OLs (PDGFRα-neg OLIG2^+^) in M1, V2 and CC of P67+42 naïve (white, n=7), P67+35CPZ+7 sham (black, n=5) or P67+35CPZ+7 iTBS (blue, n=4) mice. Repeated measures two-way ANOVA with Geisser-Greenhouse correction: treatment F (2, 13) = 22.63, p < 0.0001; region F (1.23, 15.98) = 216.1, p < 0.0001; interaction F (4, 26) = 1.533, p = 0.2219. **m.** Experimental schematic showing the time course over which *Pdgfr*α*-CreERT^TM^ :: Rosa26-YFP* mice received Tx and 28 days of LI-rTMS (sham-stimulation or iTBS). The experiment had two groups: P67+42 sham and P67+42 iTBS. **n-q.** Confocal images of YFP (green), PDGFRα (red) and OLIG2 (blue) immunohistochemistry in M1 (**n, o**) or V2 (**p, q**) of P67+42 sham (**n, p**) and P67+42 iTBS (**o, q**) mice. **r.** The proportion (%) of YFP^+^ cells that are PDGFRα-neg OLIG2^+^ OLs in the M1 and V2 cortices of P67+42 sham (n=4) and P67+42 iTBS (n=6) mice. Repeated measures two-way ANOVA: treatment F (1, 8) = 7.439, p = 0.0259; region F (1, 8) = 26.09, p = 0.0009; interaction F (1, 8) = 0.1305, p = 0.7272. Data are presented as mean ± SD. Bonferroni post- test: * p < 0.05, **** p < 0.0001, # p = 0.067. Scale bars represent 20 µm.

YFP^+^ OPCs produced new YFP^+^ OLs (OLIG2^+^ PDGFRα-neg cells) in M1 (**Fig. 1b-d**), V2 (**Fig. 1e-g**) and the CC (**Fig. 1j, k**) of P67+42 naïve and P67+35CPZ+7 sham or iTBS *Pdgfr*α*-CreERT^TM^ :: Rosa26-YFP* mice. A higher proportion of the M1 or V2 cortex YFP^+^ cells are OLs in P67+42 naïve mice relative to P67+35CPZ+7 sham or iTBS treated *Pdgfr*α*-CreERT^TM^ :: Rosa26-YFP* mice (**Fig. 1l**), indicating that the CPZ diet killed many new OLs. By contrast, the proportion of YFP^+^ cells that were new callosal OLs was equivalent in P67+42 naïve and P67+35CPZ+7 sham or iTBS treated *Pdgfr*α*-CreERT^TM^ :: Rosa26-YFP* mice (**Fig. 1l**), consistent with the rapid replacement of callosal OLs after CPZ withdrawal. Importantly, the proportion of YFP^+^ cells that were new OLs in M1, V2 or the CC was equivalent in P67+35CPZ+7 sham and iTBS mice (**Fig. 1l**), suggesting that iTBS did not alter the generation or survival of new OLs during demyelination.

However, delivering Tx to label OPCs at P60 and allowing them to generate new YFP^+^ OLs for 3 weeks (until P67+14) prior to commencing the 4-week iTBS intervention (**Fig. 1a**), ensures that only a fraction of the new YFP^+^ OLs detected at P67+42 mature under the influence of iTBS. To evaluate the impact of time-course on our conclusion, we delivered Tx to healthy P60 *Pdgfr*α*-CreERT^TM^ :: Rosa26-YFP* mice, and from P67+14 to P67+42, delivered consecutive daily sham stimulation or iTBS (**Fig. 1m**). Mice that received iTBS had ∼34% more YFP^+^ OLs in M1 and ∼41% more in V2 than sham-stimulated mice (**Fig. 1n-r**). These data are consistent with our previous report that iTBS promotes new OL survival in the mouse cortex [25], but the magnitude of the effect appears diminished when iTBS is only delivered for 4 weeks of the 7-week tracing period. This experiment allows us to be more confident in our conclusion that, unlike its effect in the healthy CNS, iTBS is unable to promote new OL survival during CPZ-feeding.

### iTBS increases the number of mGFP^+^ internodes elaborated by new M1 OLs

If iTBS could alter the number or length of myelin internodes produced by the new OLs [25], it could promote remyelination without increasing the number of new OLs added to the brain. To examine this possibility, Tx was delivered to P60 *Pdgfr*α*-CreERT^T2^ :: Tau-mGFP* mice to label OPCs and reveal the full morphology of the mGFP^+^ premyelinating and myelinating OLs they produce over time. From P67, mice received CPZ for 5 weeks before being returned to normal chow for 1 week. Mice also received sham or iTBS stimulation from P67+14 (as per **Fig. 1a**). Coronal brain sections from P67+35CPZ+7 *Pdgfr*α*-CreERT^T2^ :: Tau-mGFP* mice, that included M1, V2 or CC, were immunolabeled to detect mGFP, PDGFRα and OLIG2 (**Fig. 2a-e**). In all regions examined, mGFP^+^ OPCs (**Fig. 2a**) had differentiated to produce new mGFP^+^ premyelinating (**Fig. 2b**) and myelinating OLs (**Fig. 2c-e**). Premyelinating OLs no longer expressed PDGFRα and had a highly branched and complex morphology (**Fig. 2b**) but lacked the straight sections of mGFP^+^ labelling that denoted the myelin internodes produced by the mature and myelinating OLs (**Fig. 2c-e**).

**Figure 2:**
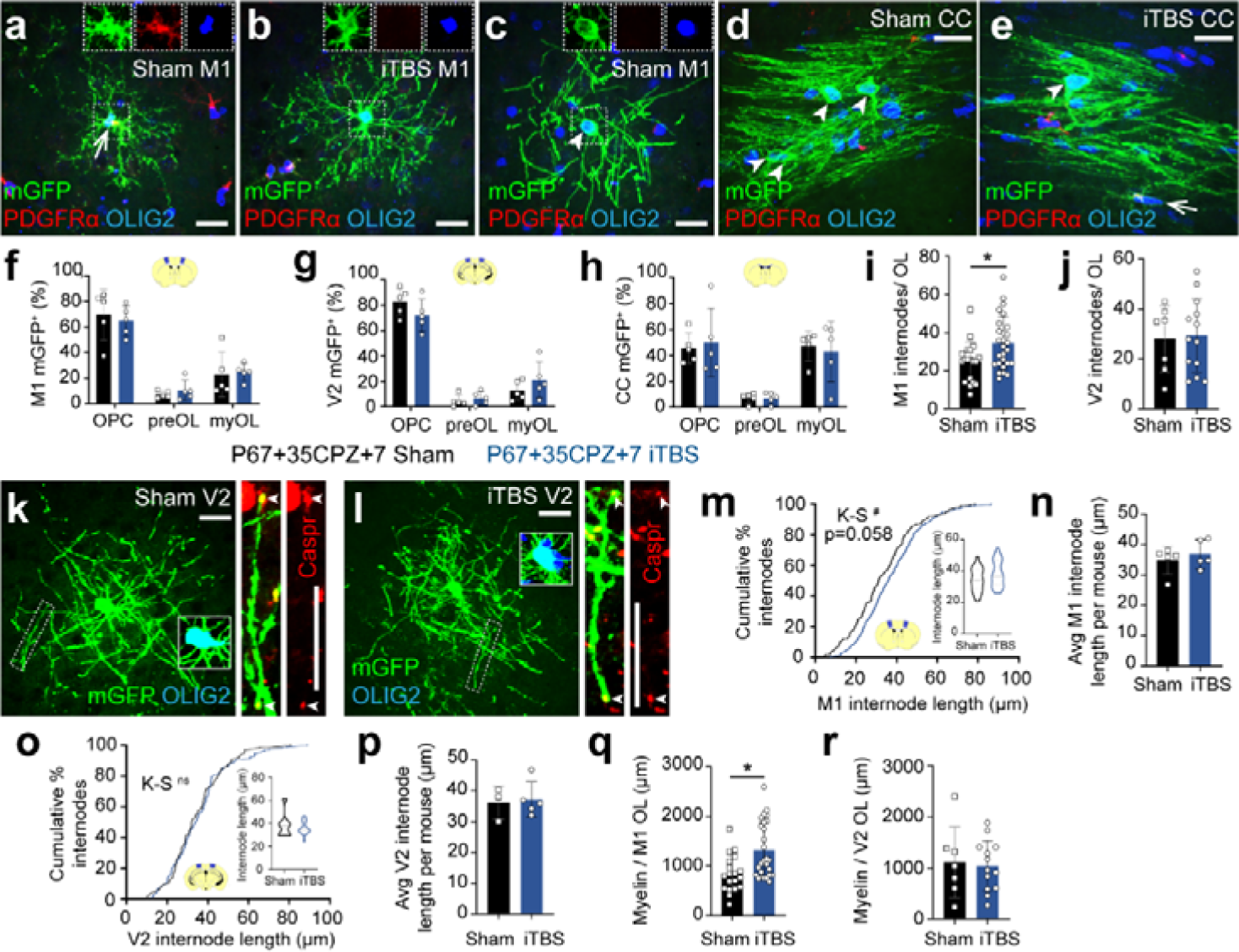
*iTBS increases the number of internodes elaborated by new OLs in M1 of CPZ-fed mice.* **a-e.** Confocal images of mGFP (green), PDGFRα (red), OLIG2 (blue) immunohistochemistry in M1 (**a-c**) and the CC (**d, e**) of *Pdgfr*α*-CreERT^T2^ :: Tau-mGFP* P67+35CPZ+7 sham (**a, c, d**) and P67+35CPZ+7 iTBS (**b, e**) mice. Arrows denote OPCs. Arrowheads denote myelinating OLs. **f-h.** The proportion (%) of mGFP^+^ cells that are OPCs, premyelinating OLs or myelinating OLs in M1 (**f**), V2 (**g**) and the CC (**h**) of P67+35CPZ+7 sham (black, n=5) and P67+35CPZ+7 iTBS (blue, n=5) mice. 95% confidence intervals overlap for P67+35CPZ+7 sham and iTBS mice, indicating no change in cell-type distribution. **i-j.** Quantification of the number of internodes produced by individual mGFP^+^ myelinating OLs in M1 (**i**) or V2 (**j**) of P67+35CPZ+7 sham or P67+35CPZ+7 iTBS mice [M1: n=19 OLs from n=5 sham mice v n=27 OLs from n=5 iTBS mice in nested t-test, p = 0.025; V2: n=7 OLs from n=3 sham mice v n=13 OLs from n=5 iTBS mice in nested t-test, p = 0.628]. **k-l.** Confocal images of mGFP (green), OLIG2 (blue; main panel insets) and CASPR (red; righthand outset) in V2 of P67+35CPZ+7 sham (**k**) or P67+35CPZ+7 iTBS (**l**) mice. The outset is a higher magnification of the region denoted by the dashed box, showing a single mGFP^+^ internode flanked by CASPR^+^ (red) paranodes (arrowheads). **m.** Cumulative distribution plot of M1 mGFP^+^ internode length for P67+35CPZ+7 sham (black) or P67+35CPZ+7 iTBS (blue) mice [n=148 sham and n=248 iTBS mGFP^+^ internodes, Kolmogorov-Smirnov (K-S) test D = 0.138, p = 0.058]. Inset violin plot for average internode length per mGFP^+^ myelinating OL in M1 of P67+35CPZ+7 sham or P67+35CPZ+7 iTBS mice [n=19 mGFP^+^ OLs from n=5 sham mice vs n=27 mGFP^+^ OLs from n=5 iTBS mice, nested t-test, p = 0.305]. **n.** Average M1 mGFP^+^ internode length per mouse for P67+35CPZ+7 sham (n = 5) and P67+35CPZ+7 iTBS (n = 5) mice [unpaired t- test, p = 0.496]. **o.** Cumulative distribution plot of V2 mGFP^+^ internode length for P67+35CPZ+7 sham (black) or P67+35CPZ+7 iTBS (blue) mice [n=66 sham and n=119 iTBS mGFP^+^ internodes, K-S test D = 0.08, p = 0.9462]. Inset violin plot of average internode length per mGFP^+^ myelinating OL in V2 of P67+35CPZ+7 sham or P67+35CPZ+7 iTBS mice [n=7 mGFP^+^ OLs from n=3 sham mice vs n=13 mGFP^+^ OLs from n=5 iTBS mice, nested t-test, p = 0.552]. **p.** Average V2 mGFP^+^ internode length per mouse for P67+35CPZ+7 sham (n=3) and P67+35CPZ+7 iTBS (n=5) mice [unpaired t-test, p = 0.79]. **q.** Estimate of myelin load per mGFP^+^ OL in M1 of P67+35CPZ+7 sham or P67+35CPZ+7 iTBS mice (avg. internode length per OL x avg. number of internodes per OL) [n=19 OLs from n=5 sham mice v n=27 OLs from n=5 iTBS mice; nested t-test, p = 0.016). **r.** Estimate of myelin load per mGFP^+^ OL in V2 of P67+35CPZ+7 sham or P67+35CPZ+7 iTBS mice (avg. internode length per OL x avg. number of internodes per OL) [n=7 OLs from n=3 sham mice v n=13 OLs from n=5 iTBS mice; nested t-test, p = 0.864]. Violin plots show the median (solid line) and interquartile range (dash lines). Data are presented as mean ± SD. * p < 0.05, # p = 0.058. Scale bars represent 20 µm.

iTBS did not alter the proportion of mGFP^+^ cells that were OPCs, premyelinating or myelinating OLs in M1, V2 or the CC (**Fig. 2f-h**). However, a more detailed analysis of the mGFP^+^ myelinating OLs revealed an increase in the average number of mGFP^+^ internodes elaborated by new M1 myelinating OLs following iTBS (∼25 internodes / OL in sham v ∼35 internodes / OL in iTBS; **Fig. 2i**). iTBS did not alter the average number of internodes elaborated by mGFP^+^ myelinating OL in V2 (**Fig. 2j**). To measure the length of mGFP^+^ internodes, coronal brain cryosections were co-labelled to detect mGFP and the contactin-associated protein (CASPR), allowing us to identify complete internodes that were flanked by CASPR^+^ paranodes (**Fig. 2k, l**). We identified a trend towards a right-ward shift in mGFP^+^ internode length distribution in M1 following iTBS (K-S test, p = 0.058, n = 148 sham and 248 iTBS internodes; **Fig. 2m**). However, average internode length per mouse was comparable in sham-stimulated and iTBS mice (**Fig. 2n**). mGFP^+^ internode length distribution and average internode length per mouse were equivalent in the V2 of P67+35CPZ+7 *Pdgfr*α*-CreERT^T2^ :: Tau-mGFP* sham- stimulated and iTBS mice (**Fig. 2o, p**). To obtain an approximation of myelin load per OL, we multiplied the average number of internodes produced per OL, with the average internode length. For sham-stimulated mice, we estimate that mGFP^+^ myelinating OLs in M1 support ∼840 µm of myelin, while those generated in iTBS mice elaborate ∼1311 µm of myelin (**Fig. 2q**). However, in V2, the estimated myelin load of mGFP^+^ myelinating OLs was equivalent between P67+35CPZ+7 sham and iTBS-treated mice (**Fig. 2r**). These data suggest that delivering iTBS during demyelination and the period immediately following CPZ withdrawal can enhance the remyelinating capacity of new myelinating OLs within M1, largely by increasing the average number of internodes elaborated per cell.

### iTBS, delivered after CPZ withdrawal, does not alter new OL number in the cortex or CC

Having established that iTBS does not increase new OL number when delivered during CPZ feeding (**Fig. 1** and **Fig. 2**), we next aimed to determine whether iTBS could increase new OL number if commenced 1 week after CPZ withdrawal (**Fig. 3**). P60 *Pdgfr*α*-CreERT^TM^ :: Rosa26-YFP* transgenic mice received Tx to label OPCs, were transferred onto a CPZ diet from P67 to induce demyelination (35 days) and then returned to a control diet to allow remyelination (**Fig. 3a**). Sham or iTBS stimulation was delivered daily for 28 days from P67+42 (7 days after CPZ withdrawal; **Fig. 3a**) and tissue collected at P67+70 for immunohistochemistry to detect YFP, PDGFRα (OPCs) and OLIG2 (cells of the OL lineage) (**Fig. 3b-g**). ∼98% of M1, ∼96% of V2, and ∼65% of CC PDGFRα^+^ OPCs were YFP-labelled in sham-stimulated P67+35CPZ+35 mice, and the labelled fraction was not significantly altered by iTBS (**Fig. 3h**). Additionally, OPC density was equivalent in sham-stimulated and iTBS mice (**Fig. 3i**). YFP^+^ PDGFRα-neg OLIG2^+^ new OLs were readily detected in the M1 (**Fig. 3b, c**), V2 (**Fig. 3d, e**) or CC (**Fig. 3f, g**) of P67+35CPZ+35 sham-stimulated or iTBS mice. Furthermore, iTBS did not alter the proportion of YFP^+^ cells that were new OLs in M1 (Sham: ∼47% of YFP^+^ OLs; iTBS: ∼50% of YFP^+^ OLs), V2 (Sham: ∼35% of YFP^+^ OLs; iTBS: ∼38% of YFP^+^ OLs) or the CC (∼72% of YFP^+^ OLs in sham and iTBS) (**Fig. 3j**).

**Figure 3:**
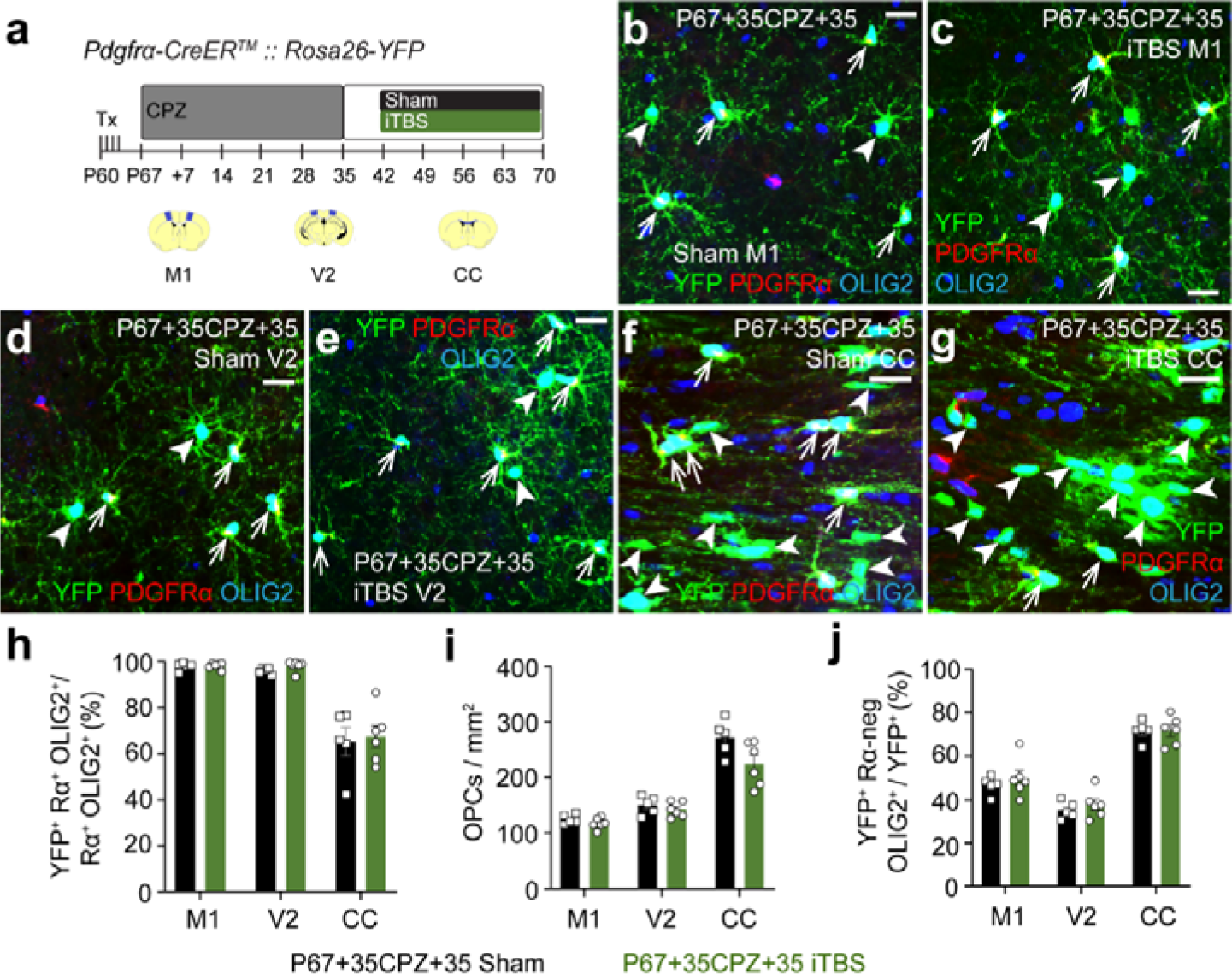
*iTBS does not increase new OL number in the cortex or CC after CPZ withdrawal.* **a.** Experimental schematic showing the timeline over which *Pdgfr*α*-CreERT^TM^ :: Rosa26-YFP* mice received 4 consecutive days of Tx gavage, 35 days of CPZ and up to 28 days of LI-rTMS (sham-stimulation or iTBS). This experiment included two treatment groups: P67+35CPZ+35 sham, and P67+35CPZ+35 iTBS. The brain regions analysed are shown in blue in the coronal brain section schematics: M1 (∼ Bregma +0.5), V2 (∼ Bregma -2.5) and the CC (∼ Bregma +0.5). **b-g.** Confocal images of YFP (green), PDGFRα (red) and OLIG2 (blue) immunohistochemistry in M1 (**b, c**), V2 (**d, e**) and CC (**f, g**) of P67+35CPZ+35 sham (**b, d, f**) or P67+35CPZ+35 iTBS (**c, e, g**) mice. Arrows indicate YFP^+^ OPCs. Arrowheads indicate new YFP^+^ OLs. **h.** The proportion (%) of PDGFRα^+^ OLIG2^+^ OPCs that are YFP-labelled in M1, V2 and the CC of P67+35CPZ+35 sham (n=5) and P67+35CPZ+35 iTBS (n=6) mice. Repeated measures two-way ANOVA with Geisser-Greenhouse correction: treatment F (1, 9) = 0.259, p = 0.623; region F (1.01, 9.11) = 75.14, p < 0.0001; interaction F (2, 18) = 0.077, p = 0.927. **i.** The density of PDGFRα^+^ OLIG2^+^ OPCs in M1, V2 and the CC of P67+35CPZ+35 sham (n=5) and P67+35CPZ+35 iTBS (n=6) mice. Repeated measures two-way ANOVA with Geisser-Greenhouse correction: treatment F (1, 9) = 5.896, p = 0.038; region F (1.221, 10.99) = 96.8, p < 0.0001; interaction F (2, 18) = 2.86, p = 0.084. **j.** The (%) of YFP+ cells that are new OLs (YFP^+^ PDGFRα-neg OLIG2^+^) in M1, V2 and the CC of P67+35CPZ+35 sham (n=5) and P67+35CPZ+35 iTBS (n=6) mice. Repeated measures two-way ANOVA with Geisser-Greenhouse correction: treatment F (1, 9) = 0.756, p = 0.407; region F (1.276, 11.48) = 96.96, p < 0.0001; interaction F (2, 18) = 0.1924, p = 0.827. Data are presented as mean ± SD. Scale bars represent 20 µm.

### iTBS increases the length of internodes elaborated by new remyelinating OLs in M1 and the CC

To determine whether iTBS alters OL maturation in remyelinating mice, we delivered Tx to P60 *Pdgfr*α*- CreERT^T2^ :: Tau-mGFP* mice to fluorescently label a subset of OPCs and trace their generation of mGFP^+^ premyelinating and myelinating OLs until P67+35CPZ+35 (**Fig. 4**). Mice received daily sham stimulation or iTBS 7 days after CPZ withdrawal (as per **Fig. 3a**). We performed immunohistochemistry and examined the morphology of individual mGFP^+^ cells to classify each as an OPC (PDGFRα^+^ OLIG2^+^), premyelinating or myelinating OL (PDGFRα-neg OLIG2^+^; **Fig. 4a, b**). Approximately 40% of the mGFP^+^ cells had differentiated into myelinating OLs in M1 (**Fig. 4c**) and ∼30% in V2 of sham-stimulated P67+35CPZ+35 mice (**Fig. 4d**) - less than the ∼61% of mGFP^+^ cells that were myelinating OLs in the CC (**Fig. 4e**), consistent with a higher rate of OL differentiation and remyelination in the CC relative to the cortex following CPZ withdrawal [6]. The proportion of mGFP^+^ cells that were premyelinating or myelinating OLs in each region was equivalent between sham and iTBS-treated P67+35CPZ+35 *Pdgfr*α*-CreERT^T2^ :: Tau-mGFP* mice (**Fig. 4c-e**).

**Figure 4:**
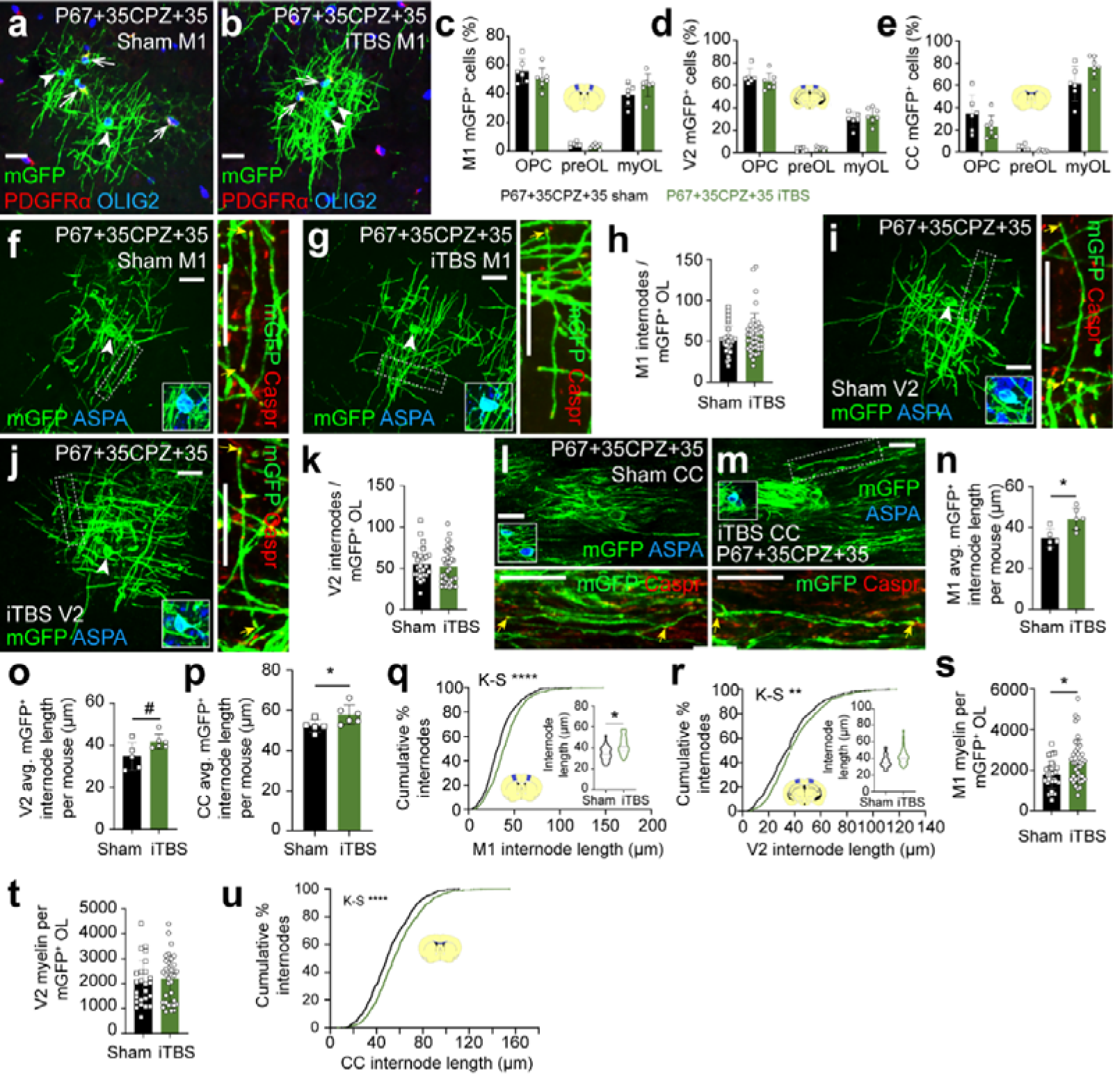
*iTBS increases the length of internodes elaborated by new OLs in M1 and the CC after CPZ withdrawal.* **a-b** Representative confocal images of mGFP (green), PDGFRα (red) and OLIG2 (blue) immunohistochemistry in M1 cortex of *Pdgfr*α*-CreERT^T2^ :: Tau-mGFP* P67+35CPZ+35 sham (**a**) and iTBS (**b**) mice. Arrows indicate mGFP^+^ OPCs. Arrowheads indicate mGFP^+^ myelinating OLs. **c-e.** The proportion (%) of mGFP cells that are OPCs, premyelinating OLs and myelinating OLs per mouse in M1 (**c**), V2 (**d**) and CC (**e**) of P67+35CPZ+35 sham (n=6) or P67+35CPZ+35 iTBS (n=7) mice. 95% confidence intervals overlap for P67+35CPZ+35 sham and P67+35CPZ+35 iTBS mice, indicating no change in cell-type distribution. **f-g.** Confocal images of mGFP (green), ASPA (blue, inset showing cell body) and CASPR (red, outset showing internode) immunohistochemistry in M1 of P67+35CPZ+35 sham (**f**) and P67+35CPZ+35 iTBS (**g**) mice. Outset shows higher magnification of dashed box with paranodes indicated by small arrows. **h.** The number of internodes elaborated by individual M1 mGFP^+^ myelinating OL in P67+35CPZ+35 sham or P67+35CPZ+35 iTBS (n=30 OLs analysed in n=5 sham mice vs n=39 OLs analysed in n=6 mice; nested t-test, p = 0.335). **i-j.** Confocal images of mGFP (green), ASPA (blue, inset showing cell body) and CASPR (red, outset showing internode) immunohistochemistry in V2 of P67+35CPZ+35 sham (**i**) and P67+35CPZ+35 iTBS (**j**) mice. Outset shows higher magnification of dashed box with paranodes indicated by small arrows. **k.** The number of internodes elaborated by individual V2 mGFP^+^ myelinating OL in P67+35CPZ+35 sham or P67+35CPZ+35 iTBS (n=25 OLs analysed in n=5 sham mice vs n=36 OLs analysed in n=6 mice; nested t-test, p = 0.731). **l-m.** Single confocal z-plane image of mGFP (green), ASPA (blue, inset showing cell body) and CASPR (red, outset showing internode) immunohistochemistry in the CC of P67+35CPZ+35 sham (**l**) and P67+35CPZ+35 iTBS (**m**) mice. Outset shows higher magnification of adjacent z-plane (l) or dashed box (m) with paranodes indicated by small arrows. **n-p.** Average mGFP^+^ internode length per mouse in M1 (**n**), V2 (**o**) or CC (**p**) of P67+35CPZ+35 sham (n=5) and P67+35CPZ+35 iTBS (n=6) mice [M1 MWU test, p = 0.012; V2 unpaired t-test, p = 0.052; CC MWU test, p = 0.03]. **q-r.** Cumulative distribution plot of M1 (**q**) or V2 (**r**) mGFP^+^ internode length for P67+35CPZ+35 sham (black) and P67+35CPZ+35 iTBS (green) mice [M1: n=538 mGFP^+^ internodes measured from n=5 sham mice vs n=713 mGFP^+^ internodes measured from n=6 iTBS mice, K-S test D = 0.184, p < 0.0001; V2: n=427 mGFP^+^ internodes measured from n=5 sham mice vs n=730 mGFP^+^ internodes measured from n=6 iTBS mice, K-S test D = 0.107, p = 0.0041]. Inset violin plots of average mGFP^+^ internode length for individual mGFP^+^ myelinating OLs in P67+35CPZ+35 sham and P67+35CPZ+35 iTBS mice [M1: n=30 mGFP^+^ OLs from n=5 sham mice vs n=39 mGFP^+^ OLs from n=6 iTBS mice, nested t-test, p = 0.0167; V2: n=25 mGFP^+^ OLs from n=5 sham mice vs n=36 mGFP^+^ OLs from n=6 iTBS mice, nested t-test, p = 0.0753]. **s-t.** Estimate of myelin load per mGFP^+^ OL in M1 (**s**) and V2 (**t**) of P67+35CPZ+35 sham or P67+35CPZ+35 iTBS mice (avg. internode length per OL x avg. number of internodes per OL) [M1: n=30 OLs from n=5 sham mice v n=39 OLs from n=6 iTBS mice, nested t-test, p = 0.0402; V2: n=25 OLs from n=5 sham mice v n=36 OLs from n=6 iTBS mice, nested t-test, p = 0.6). **u.** Cumulative distribution plot of CC mGFP^+^ internode length for P67+35CPZ+35 sham (black) and P67+35CPZ+35 iTBS (green) mice [n=632 mGFP^+^ internodes measured from n=5 sham mice vs n=680 mGFP^+^ internodes measured from n=6 iTBS mice, K-S test D = 0.107, p < 0.0001]. Data are presented as mean ± SD. Violin plots show the median (solid line) and interquartile range (dash lines). *p<0.05, **p<0.01, ****p<0.0001, ^#^p = 0.052. Scale bars represent 20 µm.

By labelling to detect mGFP, the mature OL marker, ASPA, and the paranodal marker, CASPR, we performed a more detailed analysis of the mGFP^+^ myelinating OLs in the M1 (**Fig. 4f-h**) and V2 (**Fig. 4i-k**) cortices. We found that mGFP^+^ ASPA^+^ OLs elaborate an equivalent number of internodes in sham-stimulated and iTBS mice [M1: ∼52 internodes/OL sham v ∼58 internodes/OL iTBS; V2: ∼55 internodes/OL sham v ∼52 internodes/OL in iTBS]. The high density of mGFP^+^ OLs within the remyelinating CC made it impossible to attribute individual internodes to specific new OLs, so the number of internodes per new OL was not quantified in this region (**Fig. S1**). However, in M1 (**Fig. 4f, g**), V2 (**Fig. 4i, j**) and the CC (**Fig. 4l, m**) we were able to identify mGFP^+^ internodes that were clearly flanked by CASPR^+^ paranodes and quantify new internode length (**Fig. 4n-u**). In sham-stimulated P67+35CPZ+35 *Pdgfr*α*-CreERT^T2^ :: Tau-mGFP* mice, new mGFP^+^ internodes added to the cortex were significantly shorter than those elaborated in the CC [M1 mean ∼35µm (**Fig. 4n**), V2 mean ∼35µm (**Fig. 4o**), CC mean ∼52µm (**Fig. 4p**)]. Furthermore, iTBS delivery shifted internode length distribution within M1 (**Fig. 4q**) and V2 (**Fig. 4r**) toward longer internodes, when compared with sham-stimulated mice. This corresponded to a ∼21% increase in mean internode length per mouse in M1 (**Fig. 4n**, p=0.012) and a ∼17% increase in V2 (**Fig. 4o,** p = 0.052). This equates to iTBS increasing the myelin load of M1 OLs by ∼645 µm (**Fig. 4s**). By contrast, iTBS did not significantly increase the quantity of myelin produced by new mGFP^+^ OLs in V2 (**Fig. 4t**). Within the CC, iTBS also produced a significant right-ward shift in internode length distribution, indicative of longer internodes in the CC (**Fig. 4u**), which corresponded to a ∼ 10% increase in average internode length per mouse (**Fig. 4p**). These data suggest that iTBS facilitates remyelination by enhancing internode extension by new OLs in M1 and the CC.

### Surviving OLs do not reliably express OLIG2 after 4 weeks on a CPZ diet

To fluorescently label mature OLs and map their fate during CPZ feeding, we administered Tx to P60 *Plp- CreER^T2^ :: Rosa26-YFP and Plp-CreER^T2^ :: Tau-mGFP* transgenic mice, and commenced CPZ feeding at P67 (**Fig. 5a**). To evaluate the level of demyelination achieved by P67+28CPZ, coronal brain cryosections were collected at ∼ Bregma +0.5 and used for black-gold myelin labelling (**Fig. 5b-e**). We found that the level of overt callosal demyelination was less in *Plp-CreER^T2^ :: Rosa26-YFP* mice compared with *Plp-CreER^T2^ :: Tau- mGFP* mice (**Fig. 5f**), however, CPZ-feeding did induce demyelination in both mouse cohorts.

**Figure 5:**
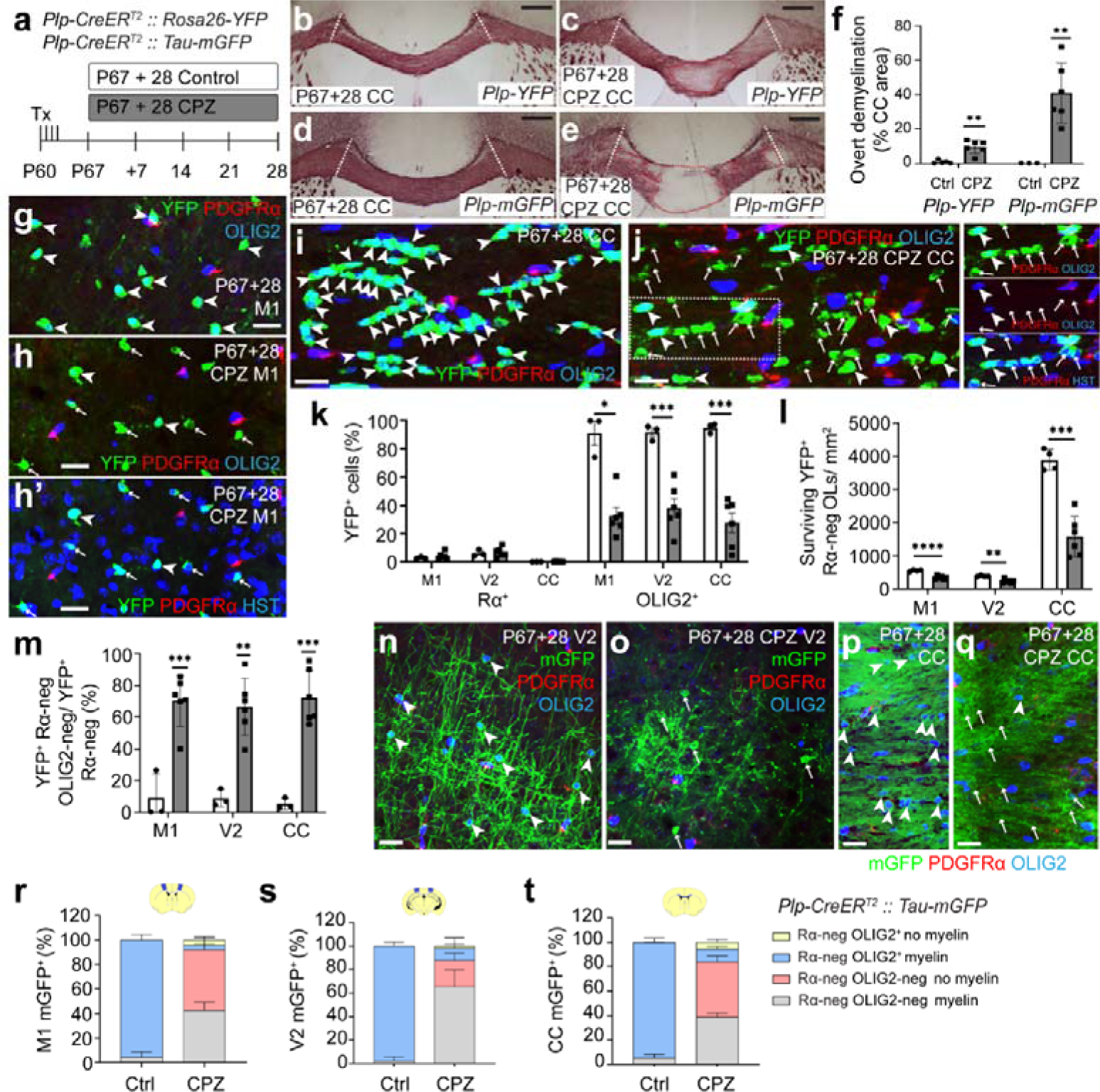
OLIG2 immunohistochemistry can become unreliable for identifying all cells of the OL lineage at 4 weeks of CPZ feeding. **a.** Experimental schematic showing the timeline over which *Plp-CreER^T2^ :: Rosa26-YFP* or *Plp-CreER^T2^ :: Tau-mGFP* transgenic mice received 4 consecutive daily doses of Tx and up to 28 days of CPZ-feeding. **b-e** Images of black-gold labelling in the CC of P67+28 control and P67+28 CPZ *Plp-CreER^T2^ :: Rosa26-YFP* mice (**b, c**), and P67+28 control and P67+28 CPZ *Plp-CreER^T2^ :: Tau-mGFP* mice (**d, e**). CC was analysed between the white dashed lines. Regions denoted by the red dashed lines show example regions of gross demyelination. **f** Area of the CC (%) where black-gold labelling was feint or absent in P67+28 control and P67+28 CPZ *Plp- CreER^T2^ :: Rosa26-YFP* mice and *Plp-CreER^T2^ :: Tau-mGFP* mice. Data from each group was evaluated using a one sample t and Wilcoxon text to determine whether the level of demyelination was different from zero [n=5 *Plp-CreER^T2^ :: Rosa26-YFP* P67+28 controls, p=0.19; n=6 *Plp-CreER^T2^ :: Rosa26-YFP* P67+28 CPZ, p=0.002; n=3 *Plp-CreER^T2^ :: Tau-mGFP* P67+28 controls (all zero); n=6 *Plp-CreER^T2^ :: Tau-mGFP* P67+28 CPZ, p=0.002]. **g-j.** Confocal images of YFP (green), PDGFRα (red), OLIG2 (blue) immunohistochemistry in M1 (**g, h**) and CC (**i, j**) of P67+28 control (**g, i**) and P67+28 CPZ (**h, j**) *Plp-CreER^T2^ :: Rosa26-YFP* mice. **h’** The same as h, however, OLIG2 labelling is substituted with Hoechst 33342 (HST, blue). Outset shows the region denoted by the dashed box as separate colour channels. Arrowheads indicate YFP^+^ PDGFRα-neg OLIG2^+^ cells. Arrows indicate YFP^+^ PDGFRα-neg OLIG2-neg cells. **k.** The proportion (%) of YFP^+^ cells that are PDGFRα^+^ OPCs or OLIG2^+^ cells of the OL lineage in M1, V2 or the CC of *Plp-CreER^T2^ :: Rosa26-YFP* P67+28 control (n=3) or *Plp-CreER^T2^ :: Rosa26-YFP* P67+28 iTBS (n=6) mice. Repeated measures two-way ANOVA with Geisser-Greenhouse correction for YFP^+^ PDGFRα^+^ [treatment F (1, 7) = 0.372, p = 0.56; region F (1.736, 12.15) = 23.67, p < 0.0001; interaction F (2, 14) = 0.24, p = 0.791] or YFP^+^ OLIG2^+^ [treatment F (1, 7) = 56.21, p = 0.0001; region F (1.282, 8.973) = 0.262, p = 0.679; interaction F (2, 14) = 0.771, p = 0.481]. **l.** The density of YFP^+^ PDGFRα-neg surviving OLs in the M1, V2 and CC of *Plp-CreER^T2^ :: Rosa26-YFP* P67+28 control (n=4) or P67+28 CPZ (n=6) mice. Repeated measures two-way ANOVA with Geisser-Greenhouse correction: treatment F (1, 8) = 48.42, p = 0.0001; region F (1.013, 8.104) = 225.2, p < 0.0001; interaction F (2, 16) = 45.84, p < 0.0001. **m.** The proportion of YFP^+^ PDGFRα-neg cells that were OLIG2-neg in M1, V2 and the CC of *Plp-CreER^T2^ :: Rosa26-YFP* P67+28 control (n=3) and P67+28 CPZ (n=6) mice. Data from each group was evaluated using a one sample t and Wilcoxon text to determine whether any group was different from the expected value of zero, followed by a Benjamini-Hochberg correction for 6 comparisons: M1 *Plp-CreER^T2^ :: Rosa26-YFP* P67+28 control, p=0.41; M1 *Plp-CreER^T2^ :: Rosa26-YFP* P67+28 CPZ, p=0.0006; V2 *Plp- CreER^T2^ :: Rosa26-YFP* P67+28 control, p=0.11; V2 *Plp-CreER^T2^ :: Rosa26-YFP* P67+28 CPZ, p=0.002; CC *Plp-CreER^T2^ :: Rosa26-YFP* P67+28 control, p=0.12; CC *Plp-CreER^T2^ :: Rosa26-YFP* P67+28 CPZ, p=0.0006. **n-q.** Confocal images of mGFP (green), PDGFRα (red) and OLIG2 (blue) immunohistochemistry in the V2 (**n, o**) or CC (**p, q**) of *Plp-CreER^T2^ :: Tau-mGFP* P67+28 control (**n, p**) and *Plp-CreER^T2^ :: Tau-mGFP* P67+28 CPZ (**o, q**) mice. Arrowheads indicate YFP^+^ PDGFRα-neg OLIG2^+^ OLs. Arrows indicate YFP^+^ PDGFRα-neg OLIG2-neg presumptive OLs. **r-t.** Stacked column charts showing the proportion of mGFP^+^ PDGFRα-neg cells that are OLIG2^+^ or OLIG2-neg with or without myelin sheaths in M1 (**r**), V2 (**s**) and the CC (**t**) of *Plp-CreER^T2^ :: Tau-mGFP* P67+28 control (n=3) and *Plp-CreER^T2^ :: Tau-mGFP* P67+28 CPZ (n=3) mice. One sample t and Wilcoxon test with Benjamini-Hochberg correction for multiple comparisons determined whether the proportion of mGFP^+^ PDGFRα-neg OLIG2-neg cells was different from zero: M1 *Plp-CreER^T2^ :: Tau-mGFP* P67+28 control, p=0.267; M1 *Plp-CreER^T2^ :: Tau-mGFP* P67+28 CPZ, p=0.003; V2 *Plp-CreER^T2^ :: Tau-mGFP* P67+28 control, p=0.423; V2 *Plp-CreER^T2^ :: Tau-mGFP* P67+28 CPZ, p=0.029; CC *Plp-CreER^T2^ :: Tau-mGFP* P67+28 Ctrl, p=0.125; CC *Plp-CreER^T2^ :: Tau-mGFP* P67+28 CPZ, p=0.001. A repeated measures two-way ANOVA with Greenhouse-Geisser correction compared the proportion of mGFP^+^ PDGFRα-neg cells without myelin sheaths: treatment F (1, 4) = 7291, p < 0.0001; region F (1.42, 5.681) = 10.1, p=0.017; interaction F (2, 8) = 10.04, p=0.007. Data are presented as mean ± SD. **p < 0.01, ***p < 0.001. Scale bars represent 100 µm (b-e) or 20 µm (g-j, n-q).

To evaluate OL survival, coronal brain cryosections were collected from *Plp-CreER^T2^ :: Rosa26-YFP* mice at ∼ Bregma +0.5 (M1 and the underlying CC) and –2.5 (V2) for immunohistochemistry to detect YFP, PDGFRα (OPCs), OLIG2 (all cells of OL lineage) or ASPA (the myelinating OL marker) and Hoechst 33342 (HST; nucleus) (**Fig. 5g-j**). A very small subset of YFP^+^ cells were PDGFRα^+^ OLIG2^+^ OPCs in M1, V2 and the CC of P67+28 control and CPZ mice (**Fig. 5k**), and in P67+28 *Plp-CreER^T2^ :: Rosa26-YFP* control mice, > 91% of YFP^+^ cells in the M1, V2 and CC were OLIG2^+^, marking them as cells of the OL lineage, largely OLs (**Fig. 5g, i, k**). In P67+28 *Plp-CreER^T2^ :: Rosa26-YFP* CPZ mice, not only was the density of YFP^+^ cells in M1, V2 and CC was significantly reduced (**Fig. 5l**), many of the YFP^+^ cells identified did not co-label for (**Fig. 5h, j, m**). These YFP^+^ PDGFRα-neg OLIG2-neg cells had a small soma with minimal cytoplasm around the nucleus, consistent with the morphology of OLs (**Fig. 5h, j**) and comprised ∼70% of M1 YFP^+^ cells, ∼67% of V2 YFP^+^ cells and ∼73% of CC YFP^+^ cells (**Fig. 5m**). As the YFP^+^ PDGFRα-neg OLIG2-neg cells were OLs prior to CPZ delivery, retained OL morphology, and the vast majority of YFP^+^ cells in M1 (∼91%) and V2 (∼96%) and ∼42% in the CC still co-labelled with the OL marker, ASPA (**Fig. S2**), we these cells are most likely surviving OLs.

A similar phenotype was detected when comparing P67+28 *Plp-CreER^T2^ :: Tau-mGFP* control and CPZ mice (**Fig. 5n-t**). In the M1, V2 and CC of P67+28 control *Plp-CreER^T2^ :: Tau-mGFP* mice, the vast majority of mGFP^+^ cells were PDGFRα-neg and OLIG2^+^ OLs and had a distinct myelinating morphology (**Fig. 5n, p, r-t**). CPZ feeding decreased the density of mGFP^+^ cells in M1 by ∼79%, V2 by ∼66% V2 and the CC by ∼66% (**Fig. S3**). However, following CPZ-feeding most of the mGFP^+^ cells remaining no longer expressed OLIG2 [∼92% of M1 mGFP^+^ cells, ∼88% of V2 mGFP^+^ cells, and ∼84% of CC mGFP^+^ cells were PDGFRα-neg OLIG2-neg cells] and only a subset continued to support mGFP^+^ internodes [∼45% of M1 mGFP^+^ cells; ∼ 75% of V2 mGFP^+^ cells and ∼50% of CC mGFP^+^ cells; **Fig. 5r-t**]. As essentially all mGFP^+^ cells in M1 and V2 of P67+28 CPZ *Plp-CreER^T2^ :: Tau-mGFP* mice co-labelled with ASPA (**Fig. S3**) and a proportion of the mGFP^+^ OLIG2- neg cells could be morphologically classified as myelinating OLs (**Fig. 5r-t**), we classified these cells as “surviving OLs”.

### iTBS delivery during CPZ feeding does not rescue pre-existing myelinating OLs

To determine whether iTBS, delivered during CPZ feeding, can alter the survival of mature OLs, Tx was administered to P60 *Plp-CreER^T2^ :: Rosa26-YFP* and *Plp-CreER^T2^ :: Tau-mGFP* transgenic mice. From P67, mice were transferred onto a CPZ diet and from P67+14 also received daily sham-stimulation or iTBS for 14 consecutive days (**Fig. 6a**). Coronal brain cryosections from P67+28CPZ *Plp-CreER^T2^ :: Rosa26-YFP* sham or iTBS mice were processed to detect YFP and ASPA (OLs) (**Fig. 6**). The density of surviving YFP^+^ ASPA^+^ OLs was equivalent in P67+28CPZ *Plp-CreER^T2^ :: Rosa26-YFP* sham and iTBS mice in M1 (Sham: ∼ 336 cells/ mm^2^, iTBS: ∼ 355 cells/ mm^2^; **Fig. 6b-c, h**), V2 (Sham: ∼ 230 cells/ mm^2^, iTBS: ∼ 263 cells/ mm^2^; **Fig. 6d-e, h**) or the CC (Sham: ∼ 1660 cells / mm^2^, iTBS: ∼ 1415 cells/ mm^2^; **Fig. 6f-g, i**). These data indicate that delivering iTBS during CPZ-feeding does not prevent OL death.

**Figure 6:**
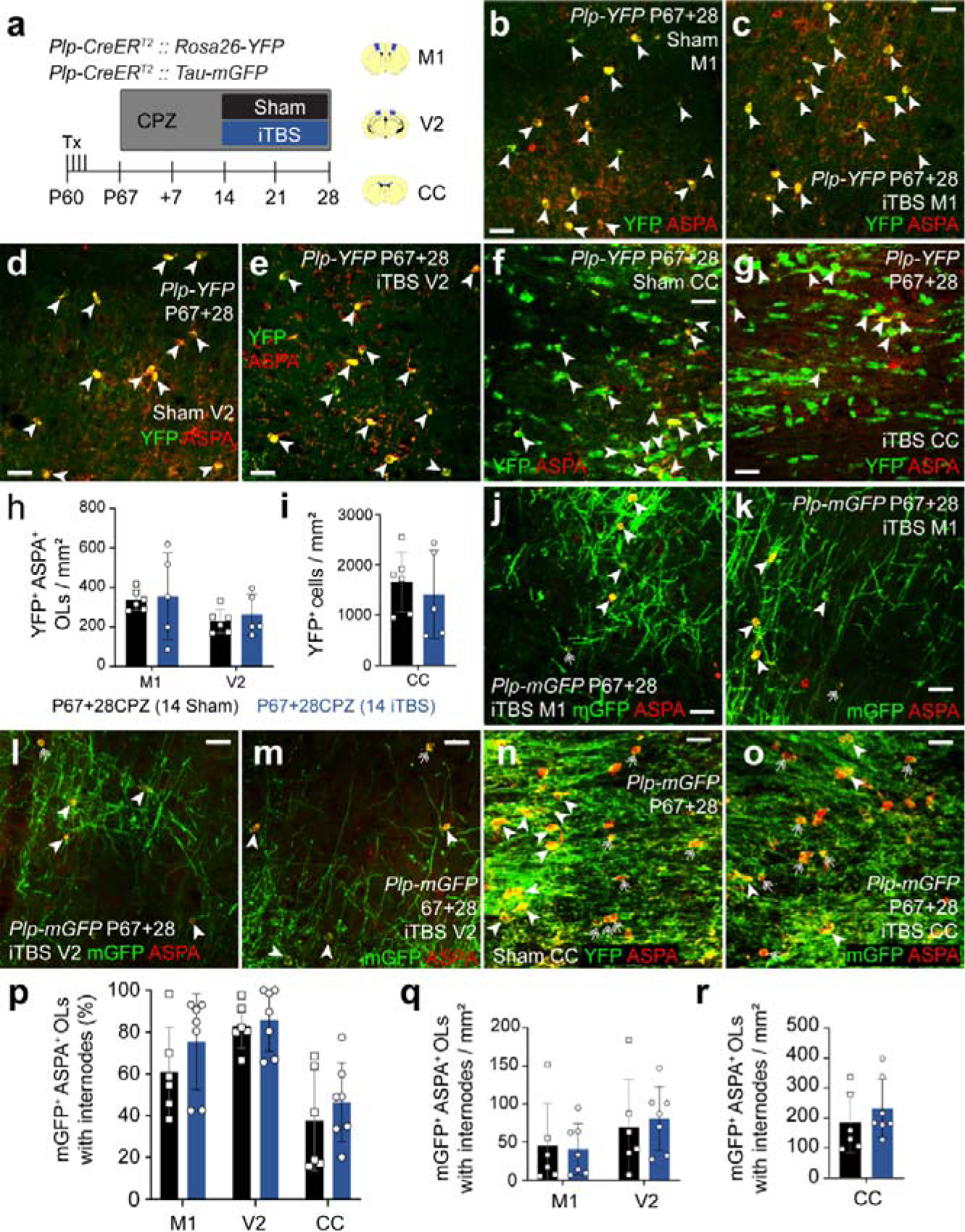
iTBS during demyelination does not affect the density of surviving and myelinating OLs in the cortex or CC. **a.** Experimental schematic showing the timeline over which *Plp-CreER^T2^ :: Rosa26-YFP* or *Plp-CreER^T2^ :: Tau-mGFP* transgenic mice received 4 consecutive daily doses of Tx, 28 days of CPZ-feeding and 14 days of LI-rTMS (sham-stimulation or iTBS). **b-g.** Confocal images of YFP (green) and ASPA (red) immunohistochemistry in M1 (**b, c**), V2 (**d, e**) or the CC (**f, g**) of *Plp-CreER^T2^ :: Rosa26-YFP* P67+28 sham or *Plp-CreER^T2^ :: Rosa26-YFP* P67+28 iTBS mice. Arrowheads indicate surviving YFP^+^ASPA^+^ OLs. **h.** The density of YFP^+^ ASPA^+^ OLs in M1 and V2 of *Plp-CreER^T2^ :: Rosa26-YFP* P67+28 sham (black, n=6) or *Plp- CreER^T2^ :: Rosa26-YFP* P67+28 iTBS (blue, n=5) mice. Repeated measures two-way ANOVA: treatment F (1, 9) = 0.146, p = 0.711; region F (1, 9) = 10.15, p = 0.011; interaction F (1, 9) = 0.045, p = 0.837. **i.** The density of YFP^+^ cells in the CC of *Plp-CreER^T2^ :: Rosa26-YFP* P67+28 sham (black, n=6) or *Plp-CreER^T2^ :: Rosa26- YFP* P67+28 iTBS (blue, n=5) mice. Unpaired t-test, p = 0.59. **j-o.** Confocal images of mGFP (green) and ASPA (red) immunohistochemistry in M1 (**j, k**), V2 (**l, m**) and the CC (**n, o**) of *Plp-CreER^T2^ :: Tau-mGFP* P67+28 sham and iTBS mice. Arrowheads indicate mGFP^+^ ASPA^+^ OLs. **p.** The proportion (%) of mGFP^+^ ASPA^+^ OLs with internodes in the M1, V2 and CC of *Plp-CreER^T2^ :: Tau-mGFP* P67+28 sham (black, n=6) and iTBS (blue, n=7) mice. Repeated measures two-way ANOVA with Geisser-Greenhouse correction: treatment F (1, 11) = 0.742, p=0.41; region F (1.753, 19.28) = 49.87, p < 0.0001; interaction F (2, 22) = 0.892, p=0.43. **q-r**. The density of mGFP^+^ ASPA^+^ OLs with internodes in M1 and V2 (**q**) or the CC (**r**) of *Plp- CreER^T2^ :: Tau-mGFP* P67+28 sham (black, n=6) and iTBS (blue, n=7) mice. Repeated measures two-way ANOVA with the Geisser-Greenhouse correction: treatment F (1, 11) = 0.29, p = 0.59; region F (1.199, 13.19) = 53.66, p < 0.0001; interaction F (2, 22) = 1.262, p = 0.303. Data are presented as mean ± SD. Scale bars represent 20 µm.

CPZ can induce a dying-back process, whereby some of the myelin sheaths can degenerate while the cell bodies remain [41, 42]. To determine whether iTBS affects the number of myelin sheaths that surviving OLs support, we examined the morphology of mGFP^+^ ASPA^+^ OLs within M1, V2 and the CC of P67+28CPZ *Plp-CreER^T2^ :: Tau-mGFP* sham and iTBS mice (**Fig. 6j-o**). In the cortex of P67+28CPZ *Plp-CreER^T2^ :: Tau-mGFP* sham mice, most of the surviving mGFP^+^ ASPA^+^ OLs retained some internodes (sham stimulated: M1 ∼13 OLs without internodes / mm^2^ vs. ∼45 OLs with internodes / mm^2^; V2 ∼10 OLs without internodes / mm^2^ vs ∼70 OLs with internodes / mm^2^; **Fig. S4**). By contrast, within the CC, most mGFP^+^ ASPA^+^ OLs lacked internodes (sham stimulated: ∼364 OLs without internodes / mm^2^ vs. ∼184 OLs with internodes / mm^2^; **Fig. S4**). Importantly, we found that the proportion of mGFP^+^ ASPA^+^ OLs that elaborated internodes in M1, V2 or the CC was equivalent in P67+28CPZ *Plp-CreER^T2^ :: Tau-mGFP* sham and iTBS mice (**Fig. 6p**). Similarly, the density of surviving OLs that retained internodes was equivalent in P67+28CPZ *Plp-CreER^T2^ :: Tau-mGFP* sham and iTBS mice (**Fig. 6j-o, q, r**). These data indicate that 4 weeks of CPZ feeding results in a higher proportion of mature OLs losing their myelin internodes in the CC relative to the cortex, but also indicate that delivering iTBS during CPZ-feeding does not affect the number of “bare” OLs.

### iTBS increases the contribution that surviving OLs make to remyelination in the CC

As iTBS can increase the myelin load of new M1 OLs during and after CPZ feeding, it may also support internode elaboration by the OLs that survive a demyelinating event [10, 41]. To determine whether iTBS can increase the contribution that surviving OLs make to myelin repair, a second cohort of *Plp-CreER^T2^ :: Rosa26- YFP* and *Plp-CreER^T2^ :: Tau-mGFP* mice commenced CPZ feeding at P67, but were returned to normal chow from P67+35. One week later, they received their first of 28 consecutive daily sham or iTBS sessions (**Fig. 7a**). Coronal brain cryosections from P67+35CPZ+35 *Plp-CreER^T2^ :: Rosa26-YFP* (**Fig. 7b-i**) and *Plp-CreER^T2^ :: Tau-mGFP* mice (**Fig. 7j-r**) were processed to detect YFP or mGFP and ASPA. By P67+35CPZ+35, essentially all YFP^+^ cells express ASPA, identifying them as surviving OLs. We note that the density of YFP^+^ ASPA^+^ OLs in M1 fell significantly between P67+28CPZ and P67+35CPZ+35 (compare M1 sham data in **Fig. 6h** and **Fig. 7h** or see **Fig. S4**, p = 0.04) but was unchanged in V2 or the CC (compare **Fig. 6h, i** with **Fig. 7h, i** or see **Fig. S4**, p > 0.99). These data suggest that mature OLs die in M1 over a longer time-course than those in V2 or the CC. However, as the density of YFP^+^ ASPA^+^ OLs in M1, V2 or the CC was equivalent in P67+35CPZ+35 *Plp-CreER^T2^ :: Rosa26-YFP* sham and iTBS mice, iTBS does not modify the survival of mature OL after CPZ withdrawal (**Fig. 7h, i**).

**Figure 7:**
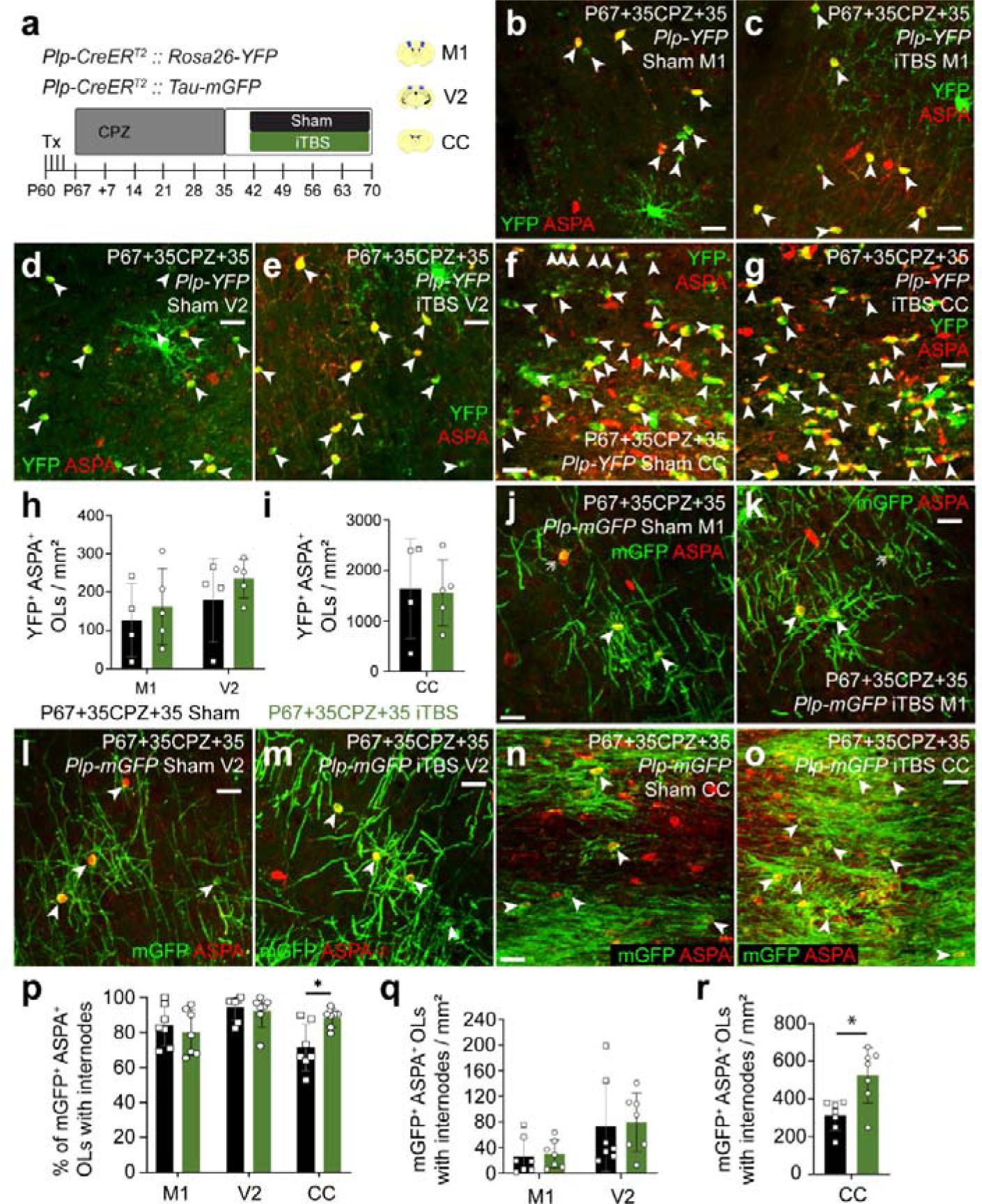
iTBS during remyelination increases the number of surviving myelinating OLs in the corpus callosum. **a.** Experimental schematic showing the timeline over which *Plp-CreER^T2^ :: Rosa26-YFP* or *Plp-CreER^T2^ :: Tau-mGFP* transgenic mice received 4 consecutive daily doses of Tx, 35 days of CPZ-feeding and 28 days of LI-rTMS (sham-stimulation or iTBS). **b-g.** Confocal images of YFP (green) and ASPA (red) immunohistochemistry in M1 (**b, c**), V2 (**d, e**) or the CC (**f, g**) of *Plp-CreER^T2^ :: Rosa26-YFP* P67+35CPZ+35 sham (**b, d, f**) or iTBS (**c, e, g**) mice. Arrowheads indicate YFP^+^ ASPA^+^ OLs. **h-i.** The density of YFP^+^ ASPA^+^ OLs in M1 and V2 (**h**) or the CC (**i**) of *Plp-CreER^T2^ :: Rosa26-YFP* P67+35CPZ+35 sham (black, n=4) or iTBS (green, n=5) mice. Repeated measures two-way ANOVA with Geisser-Greenhouse correction: treatment F (1, 7) = 0.0003, p=0.986; region F (1.011, 7.074) = 32.88, p=0.0007; interaction F (2, 14) = 0.066, p=0.936. **j-o.** Confocal images of mGFP (green) and ASPA (red) immunohistochemistry in M1 (**j, k**), V2 (**l, m**) or the CC (**n, o**) of *Plp-CreER^T2^ :: Tau-mGFP* P67+35CPZ+35 sham or iTBS mice. Arrowheads indicate mGFP^+^ ASPA^+^ OLs. **p.** The proportion (%) of mGFP^+^ ASPA^+^ OLs with internodes in M1, V2 or the CC of *Plp-CreER^T2^ :: Tau-mGFP* P67+35CPZ+35 sham (black, n=7) or iTBS (green, n=7) mice. Repeated measures two-way ANOVA with Geisser-Greenhouse correction: treatment F (1, 12) = 0.884, p = 0.366; region F (1.577, 18.92) = 8.652, p = 0.004; interaction F (2, 24) = 5.613, p = 0.01. **q-r.** Quantification of the density of mGFP^+^ ASPA^+^ OLs with internodes in M1 and V2 (**q**) or the CC (**r**) of *Plp-CreER^T2^ :: Tau-mGFP* P67+35CPZ+35 sham (black, n=7) or iTBS (green, n=7) mice. Repeated measures two-way ANOVA with Geisser-Greenhouse correction: treatment F (1, 12) = 6.546, p = 0.025; region F (1.148, 13.77) = 135.5, p < 0.0001; interaction F (2, 24) = 10.76, p = 0.0005. Data are presented as mean ± SD. *p < 0.05. Scale bars represent 20 µm.

By analysing the morphology of mGFP^+^ ASPA^+^ OLs in sham-stimulated *Plp-CreER^T2^ :: Tau-mGFP* mice, we found that the proportion of OLs with internodes did not change significantly in M1 or V2 between P67+28CPZ and P67+35CPZ+35 (**Fig. S4**). By contrast, more surviving mGFP^+^ ASPA^+^ callosal OLs support internodes in the CC of P67+35CPZ+35 mice relative to P67+28CPZ mice (compare sham data in **Fig. 6p, r** with **Fig. 7p, r** or see **Fig. S4**). This equates to ∼184 mGFP^+^ ASPA^+^ OLs with internodes / mm^2^ at P67+28CPZ vs. ∼313 mGFP^+^ OLs with internodes/ mm^2^ at P67+35CPZ+35. These data confirm that at least a subset of surviving callosal OLs spontaneously regrow their myelin sheaths upon CPZ withdrawal. We found that delivering iTBS did not alter the proportion or density of mGFP^+^ OLs that elaborated internodes in M1 or V2 of P67+35CPZ+35 *Plp-CreER^T2^ :: Tau-mGFP* mice (**Fig. 7p, q**). However, iTBS significantly increased the proportion of mGFP^+^ OLs that had internodes (**Fig. 7p**). This increased the density of myelinating mGFP^+^ OLs by ∼41%, relative to sham-stimulated mice (**Fig. 7r**). These data indicate that iTBS, delivered during remyelination, can significantly increase the number of surviving callosal OLs that contribute to remyelination.

## Discussion

LI-rTMS, applied in an iTBS pattern, modifies internode extension by new and surviving OLs to enhance remyelination. When iTBS was delivered during CPZ feeding, it did not increase new OL number (**Fig. 1**) or enhance the survival of myelinating OLs (**Fig. 6**) but increased the number of internodes elaborated by new M1 OLs (**Fig. 2**). This effectively increased the myelin load of each new M1 OL by ∼471µm. When iTBS was delivered after CPZ withdrawal (during remyelination), it did not increase the addition of new OLs to M1, V2 or the CC (**Fig. 3**). Instead, iTBS significantly increased the length of the internodes produced by new M1 and callosal OLs (**Fig. 4**) and increased the density of surviving callosal OLs that contributed to remyelination (**Fig. 7**). While the effect of LI-rTMS may be small at the level of individual internodes, its overall effect on remyelination would be substantial, indicating that this non-invasive intervention may be effective at promoting remyelination in the context of demyelinating diseases such as MS.

### iTBS does not promote new OL survival during or after CPZ feeding

When LI-rTMS is delivered during CPZ feeding (**Fig. 1**) or after CPZ withdrawal (**Fig. 3**), an equivalent number of new OLs were detected in the M1 or V2 cortices of sham-stimulated and iTBS mice. ∼85% of new M1 OLs produced during CPZ feeding die [41], so our data indicate, perhaps unsurprisingly, that iTBS cannot save new OLs from toxin-mediated cell death. However, even after CPZ withdrawal, iTBS did not alter new OL number in any of the brain regions examined (**Fig. 3**). By commencing iTBS 21 or 49 days after Tx administration, our quantification of new YFP^+^ OLs inevitably included new OLs that matured under equivalent conditions in sham and iTBS mice, prior to the LI-rTMS period. This diluted our capacity to see any effect of iTBS on new OL number. Indeed, our capacity to discern the effect of iTBS on new OL survival in the cortex of healthy mice was diminished by commencing stimulation 21 days after Tx delivery (**Fig. 1**) instead of 7 days after Tx delivery [25], and would be further diminished by a 49-day delay (as in **Fig. 3**). Therefore, we can conclude that when delivered during CPZ demyelination or remyelination, iTBS does not exert a gross effect on new OL number, but we cannot rule out some small effect on cell survival in this or other demyelinating contexts.

### iTBS increases the myelin load of new OLs in M1, but not V2

Grey matter demyelination can be extensive in the brains of people with MS, often exceeding the more studied white matter demyelination [43, 44]. A high cortical lesion load is associated with physical disability and cognitive dysfunction in MS [45, 46], making cortical remyelination an important therapeutic objective for MS research. By performing transgenic lineage tracing to identify OLs produced from parenchymal OPCs in response to the injury, we were able to study their morphology. When iTBS was delivered during CPZ feeding and for the first 7 days following CPZ withdrawal, it increased the number of internodes elaborated by new M1 OLs (**Fig. 2**). When we instead started iTBS 7 days after CPZ withdrawal, it had no effect on the number of internodes elaborated by new M1 OLs (**Fig. 4**). Instead, it increased the length or cumulative length distribution of internodes elaborated by new M1, V2 and CC OLs (**Fig. 4**). This represented a small increase in internode length, as internodes were, on average, ∼8 µm longer for new M1 OLs, however, this corresponded to ∼645 µm more myelin per M1 OL. When considered in the context of the large number of new internodes elaborated in each region, this represents a substantial increase in remyelination.

It is unclear why iTBS is more effective at promoting remyelination by new OLs in the M1 than V2 cortex, when it can equally effect OLs in these regions of the uninjured brain [25]. This regional difference could result from regional differences in the level of OL death and severity of demyelination, or cortical organisation and the way the neurons respond to demyelination. CPZ does not produce a consistent level of demyelination along the rostro-caudal axis of the CC, with the rostral region generally experiencing less demyelination than the caudal region [36, 37]. It is unclear whether this is also true in the cortex, however, we found that 4 weeks of CPZ feeding killed ∼39% of the YFP^+^ PDGFRα-neg OLs in the M1 and ∼40% in the V2 of *Plp-CreER^T2^ :: Rosa26- YFP* mice (**Fig. 5**), indicating the initial loss of mature OLs induced by CPZ was equivalent in M1 and V2 cortices. However, we found that OL loss continued between P67+28CPZ and P67+35CPZ+35 mice, consistent with a previous study of CPZ-induced OL death in M1 [41], but OL density remained stable in V2 (**Fig. S4**). Additionally, after 4 weeks of CPZ feeding, only ∼46% of surviving M1 OLs retained internodes, while ∼76% of V2 OLs elaborated internodes (**Fig. 5**). These data suggest that the pattern of OL loss and demyelination is different in M1 and V2 which, on top of structural differences between the M1 and V2 cortical circuitry [47], could influence the way that cells respond to iTBS.

The outcome of rTMS is significantly influenced by the baseline neuronal activity of the stimulated region [48]. It is possible that the baseline neuronal activity of M1 and V2 change significantly following demyelination. Multisite electrode recordings of the extracellular potentials of neurons in the motor cortex, indicate that median neuronal firing rate increases by ∼70% in the week following CPZ-withdrawal, but returns to normal by 3 weeks [41]. This had a significant impact on the outcome of a motor learning intervention, which was ineffective during the period of hyperactivity but increased the proportion of myelin sheathes that faithfully replaced lost internodes, if delivered after activity returned to normal [41]. The transient increase in neuronal firing in the motor cortex that subsides with remyelination may also explain why iTBS delivered until 1 week of remyelination is associated with new M1 OLs producing more internodes, while iTBS delivered from 1 week of remyelination is instead associated with longer internodes. Not only are regional and temporal differences in baseline GABAergic or glutamatergic signalling likely to influence the outcome of iTBS, but iTBS can work by modulating neurotransmitter release, and changes in GABAergic and glutamatergic signalling can directly influence internode length [49, 50].

### iTBS can promote remyelination by increasing myelin production by surviving OLs

A higher proportion of surviving OLs myelinate axons in the CC of P67+35CPZ+35 iTBS compared with sham- stimulated mice (**Fig. 7**). This could be the result of iTBS preventing internode loss or supporting internode regeneration. As the density of surviving callosal OLs that myelinate is significantly lower at 4 weeks of CPZ- feeding (P67+28CPZ) than after 5 weeks of recovery (P67+35CPZ+35) (**Fig. S4**), it can only be explained by the later. Internode preservation could maintain the density of myelinating OLs over time but cannot account for an increase.

Researchers have taken a variety of experimental approaches to gather evidence supporting the idea that OLs can survive a demyelinating injury and regenerate internodes [7, 10, 41, 51, 52]. In a feline model of spinal cord demyelination, the g-ratio measured for axons myelinated by individual ventral column OLs becomes highly variable, suggesting the OLs support mature and remyelinating sheathes [10]. The birth-dating of OLs within MS shadow-plaques, which are traditionally considered partially remyelinated lesions, also suggested that surviving, developmentally generated OLs contribute to human remyelination [7]. However, definitive evidence of surviving OL being able to grow new internodes came from live imaging studies that followed the fate of OLs during demyelination and remyelination in the zebrafish spinal cord or mouse cortex [41, 51, 53]. In the zebrafish spinal cord, surviving OLs generated a small number of new internodes, but these frequently and inappropriately myelinated neuronal cell bodies [51]. In the mouse cortex, 3 weeks after CPZ-withdrawal, surviving OLs were more likely to remyelinate the denuded axons than new OLs [41]. But under more inflammatory conditions, only a very small proportion of surviving cortical OLs elaborate new internodes [53]. Perhaps unsurprisingly, clemastine, which acts on OPCs to enhance remyelination, did not increase remyelination by surviving OLs [53]. There is some evidence that the sustained activation of extracellular signal-regulated kinases 1 and 2 (ERK1/2) can increase the number of surviving OLs that ensheathe demyelinated axons [52]. Furthermore, a motor learning intervention 10-17 days after cuprizone withdrawal, can dramatically increase the proportion of surviving OL that make new myelin sheathes [41]. However, more research is required to determine how surviving OLs mechanistically extend new processes and wrap new sheathes, and how this is supported by iTBS delivery during remyelination.

### The potential use of LI-rTMS as a remyelinating therapy for people with MS

LI-rTMS may be useful as an adjunct to immune modulatory therapy for the treatment of MS. Drugs or interventions that increase remyelination remain an unmet need for people with MS, as they hold the potential to restore neuron function, promote neuroprotection by limiting neurodegeneration, and increase functional recovery [54, 55]. We report that LI-rTMS, delivered in an iTBS pattern, can increase remyelination following CPZ-induced demyelination, by affecting the behaviour of new and surviving OLs. However, the outcome may depend on the timing of LI-rTMS delivery relative to lesion development, and the level of inflammation within the CNS. It was recently shown that surviving OLs in the inflamed and demyelinated cortex, elaborate a very small number of internodes and these are shorter than those measured under control conditions [53]. The CPZ- model of demyelination is associated with increased astrogliosis and microgliosis [56] but is not a model of inflammatory demyelination. Therefore, it would be interesting to determine whether LI-rTMS can improve the ability of surviving OLs to contribute to repair under more overtly inflammatory conditions. In people with MS, immune modulatory drugs can reduce the frequency of clinical relapses and impede peripheral immune cell egress into the CNS, however, treatments are still required that will prevent or reverse neuroinflammation, in the form of astrogliosis and microgliosis, which is associated with more severe disease progression [57, 58]. Several lines of evidence indicate that rTMS can direct astrocytes and microglia to transition from a pro- inflammatory to an anti-inflammatory phenotype when delivered following a CNS injury [59–65]. LI-rTMS exerts a cell autonomous effect on astrocytes, as a single session delivering 600 pulses (1 or 10 Hz, 18mT) downregulates the expression proinflammatory genes in cultured mouse primary astrocytes [66]. The capacity for LI-rTMS to jointly suppress neuroinflammation and promote remyelination, suggests that it could be beneficial for people with MS on multiple fronts, however our studies also suggest that care will need to be taken to time LI-rTMS delivery to maximise its ability of to promote myelin repair, and this could fluctuate across the disease course.

### Considerations when using immunohistochemistry to quantify OL loss in mouse models of demyelination

The basic helix-loop-helix transcription factor, OLIG2, is expressed by cells of OL lineage [67] and while it is a reliable marker for cells of the OL lineage in healthy adult mice, our data suggest that there are brief windows during CPZ-feeding where OLIG2 labelling is poor and its use to identify surviving OLs, misleading. By performing lineage tracing of mature OLs during CPZ feeding, we determined that after 4 weeks of CPZ feeding, a significant proportion of OLs within the cortex or CC cannot be identified by OLIG2 labelling alone (**Fig. 5**). The issue with OLIG2 immunohistochemistry was restricted to pre-existing OLs, as OLIG2 was clearly visible in PDGFRα^+^ OPCs in these regions. It has been reported that the OL soma can remain intact after demyelination – this has been described as dying-back oligodendrogliopathy [42]. Some of the surviving OLs that lacked OLIG2 had no remaining internodes while others supported some myelin internodes, and OLIG2 expression was recovered by all surviving OLs following CPZ withdrawal. We find this observation noteworthy, because, for a time, if these OLs had not been fluorescently labelled by our transgenic lineage tracing approach, they would have been “invisible” based on OLIG2 labelling and incorrectly assumed to have died. The transient loss of OLIG2 immunohistochemistry may be functionally significant, as OLIG2 regulates key aspects of OL development, including differentiation and myelination [67, 68] and the genetic ablation of OLIG2 increases p53 activation leading to neural progenitor and OPC death [69]. As the activation of p53 can mediate OL apoptosis in CPZ-fed mice [70], it is possible that reduced OLIG2 expression precedes the p53- dependent apoptotic death of OLs in this model. However, this is certainly not true for all OLIG2-neg OLs, as many survived and recovered their OLIG2 labelling. Our data suggest that ASPA labelling detects a higher proportion of the surviving OLs than OLIG2 during CPZ-feeding and indicate that OL loss may be dramatically overestimated in studies that solely rely on immunohistochemical readouts.

## Author contributions

P.T.N, M.C, K.M, T.S.L, L.R., C.L.C and K.M.Y carried out the experiments and / or quantification. P.T.N and K.M performed the statistical analyses. K.M.Y., J.R., C.L.C., K.M. and P.T.N designed the experiments. K.M.Y, B.V.T and K.M. provided supervision. K.M.Y., C.L.C., B.V.T., and J.R. secured project funding. P.T.N and K.M.Y wrote the manuscript. K.M., J.R., B.V.T. and C.L.C edited the manuscript.

## Data availability

The image files and data described in this manuscript can be obtained from the corresponding author by reasonable request.

## Ethics approval

All animal experiments were carried out on project A0018606, which was approved by the University of Tasmania’s Animal Ethics Committee.

## Consent for publication

This publication does not include data from human research participants.

