## Supplementary Figures for "Low intensity repetitive transcranial magnetic stimulation enhances remyelination by newborn and surviving oligodendrocytes in the cuprizone model of toxic demyelination"

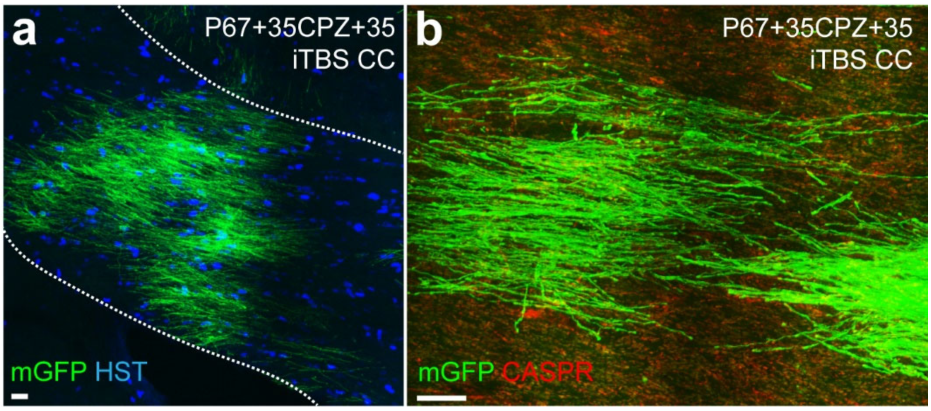


***Figure S1: High density of mGFP⁺ myelin internodes were generated by new myelinating oligodendrocytes in the CC of Pdgfrα-CreERT$^{T2}$ :: Tau-mGFP mice.***

**a-b** Compressed confocal images of the CC of a P67 *Pdgfrα-CreERT$^{T2}$ :: Tau-mGFP* mouse that had been administered 4 doses of Tx at P60 before they received 35 days of CPZ followed by 35 days of CPZ cessation and commenced 28 days of iTBS from 7 days of CPZ withdrawal. This section was immunolabeled with mGFP and HST to visualise new myelinating mGFP$^+$ OLs under 20 × objective (**a**) and immunolabeled with mGFP and CASPR to visualise individual mGFP$^+$ internodes flanked by CASPR$^+$ paranodes under 40 × objective (**b**). It was not possible to discern individual mGFP$^+$ newly generated OLs and their associated myelin internodes within the CC. Scale bars represent 20 µm.

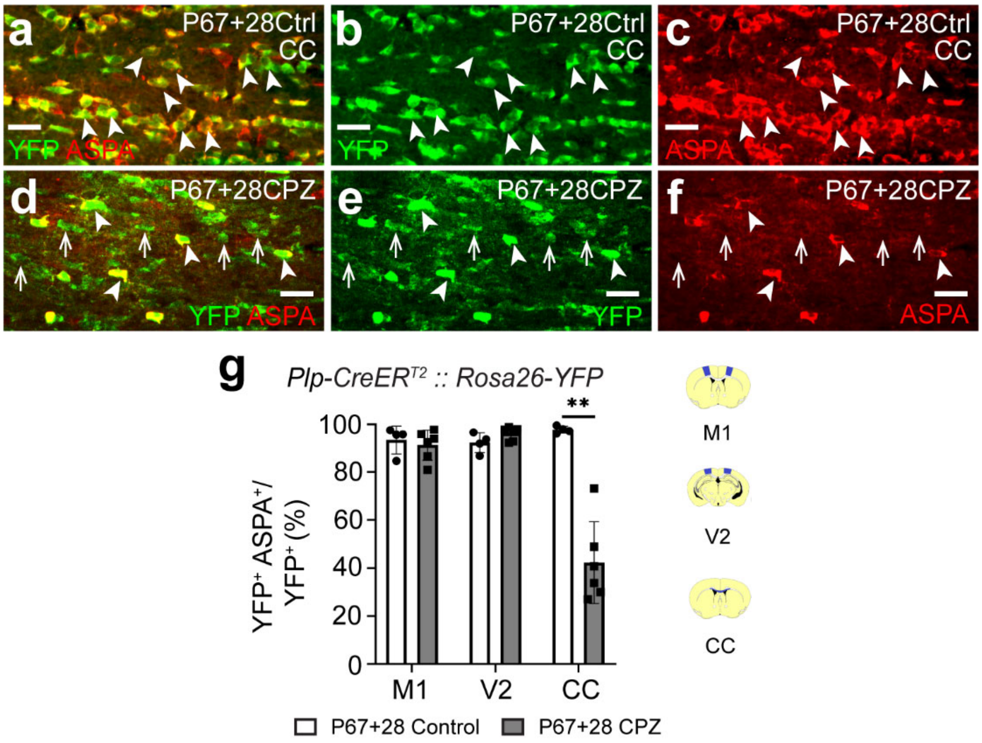


***Figure S2: A high proportion of $YFP^+$ cells no longer co-express ASPA in the CC of P67+28 CPZ Plp-CreER$^{T2}$ :: Rosa26-YFP mice.***

**a-f.** Confocal images of YFP (green) and ASPA (red) immunohistochemistry in the CC of P67+28 Control (**a-c**) and P27+28 CPZ (**d-f**) *Plp-CreER$^{T2}$ :: Rosa26-YFP* mice. Arrowheads denote $YFP^+$ $ASPA^+$ OLs. Arrows denote $YFP^+$ cells that do not express ASPA. **g.** The proportion (%) of $YFP^+$ cells that co-express ASPA in M1, V2 and the CC of P67+28 Control *Plp-CreER$^{T2}$ :: Rosa26-YFP* mice (n=4) or P67+28 CPZ *Plp-CreER$^{T2}$ :: Rosa26-YFP* mice (n=6). Repeated measures two-way ANOVA with Geisser-Greenhouse correction: treatment $F (1, 8) = 29.3$, $p = 0.0006$; region $F (1.429, 11.43) = 22.65$, $p = 0.0002$; interaction $F (2, 16) = 33.33$, $p < 0.001$. Schematics of coronal brain sections show the analysed regions in blue i.e. M1 (~ Bregma +0.5), V2 (~ Bregma -2.5) and CC underlying M1 (~ Bregma +0.5). Data are presented as mean ± SD. Bonferroni post-test: $^{**}p < 0.01$. Scale bars represent 20 µm.

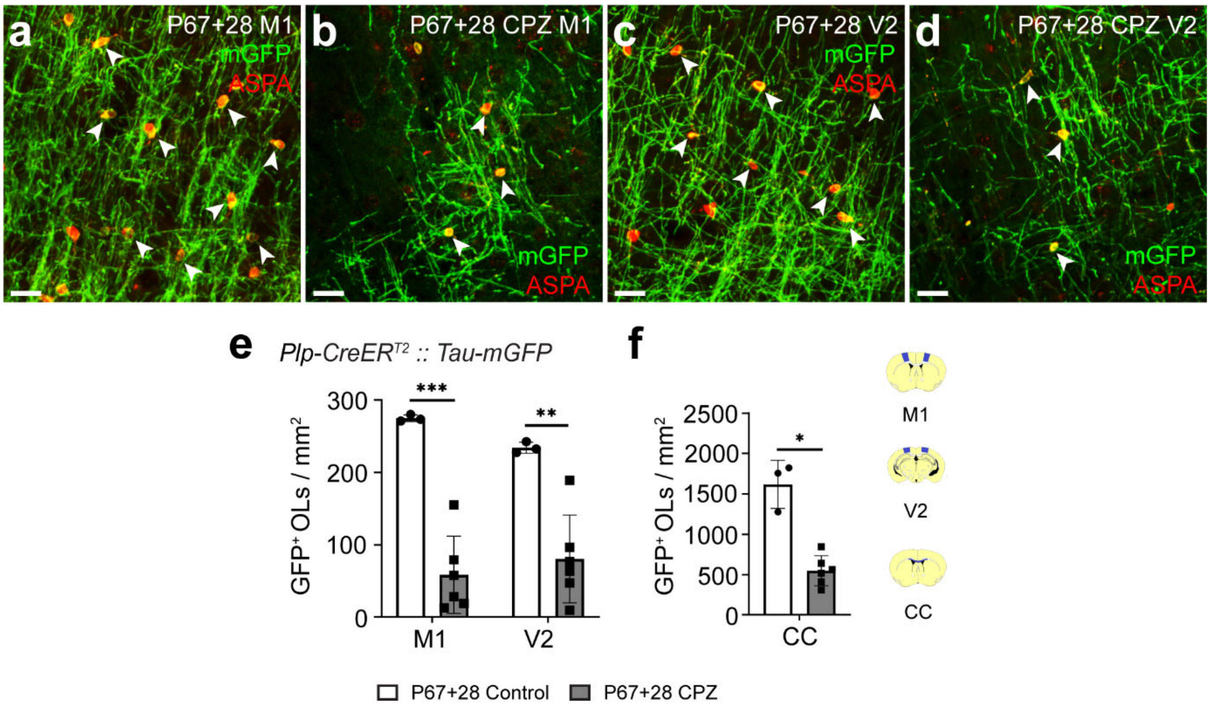


***Figure S3: CPZ-feeding reduces the density of mGFP$^+$ OLs in the M1, V2 and CC of Plp-CreER$^{T2}$ :: Tau-mGFP mice.***

**a-d** Confocal images of mGFP (green) and ASPA (red) immunohistochemistry in M1 (**a, b**) and V2 (**c, d**) of *Plp-CreER$^{T2}$ :: Tau-mGFP* P67+28 control (**a, c**) and *Plp-CreER$^{T2}$ :: Tau-mGFP* P67+28 CPZ (**b, d**) mice. Arrowheads indicate mGFP$^+$ ASPA$^+$ OLs. **e-f** The density of mGFP$^+$ cells (OLs) in M1, V2 (**e**) and the CC (**f**) of P67+28 Control *Plp-CreER$^{T2}$ :: Tau-mGFP mice* (n=3) and P67+28CPZ *Plp-CreER$^{T}$ :: Tau-mGFP* mice (n=6). Repeated measures two-way ANOVA with Geisser-Greenhouse correction: treatment $F(1, 7) = 79.68$, $p < 0.0001$; region $F(1.215, 8.505) = 120.7$, $p < 0.0001$; interaction $F(2, 14) = 27.87$, $p < 0.0001$. Schematics of coronal brain sections show the analysed regions in blue i.e. M1 (~ Bregma +0.5), V2 (~ Bregma -2.5) and CC underlying M1 (~ Bregma +0.5). Data are presented as mean ± SD. Bonferroni post-test: *$p < 0.05$, **$p < 0.01$, ***$p < 0.001$.

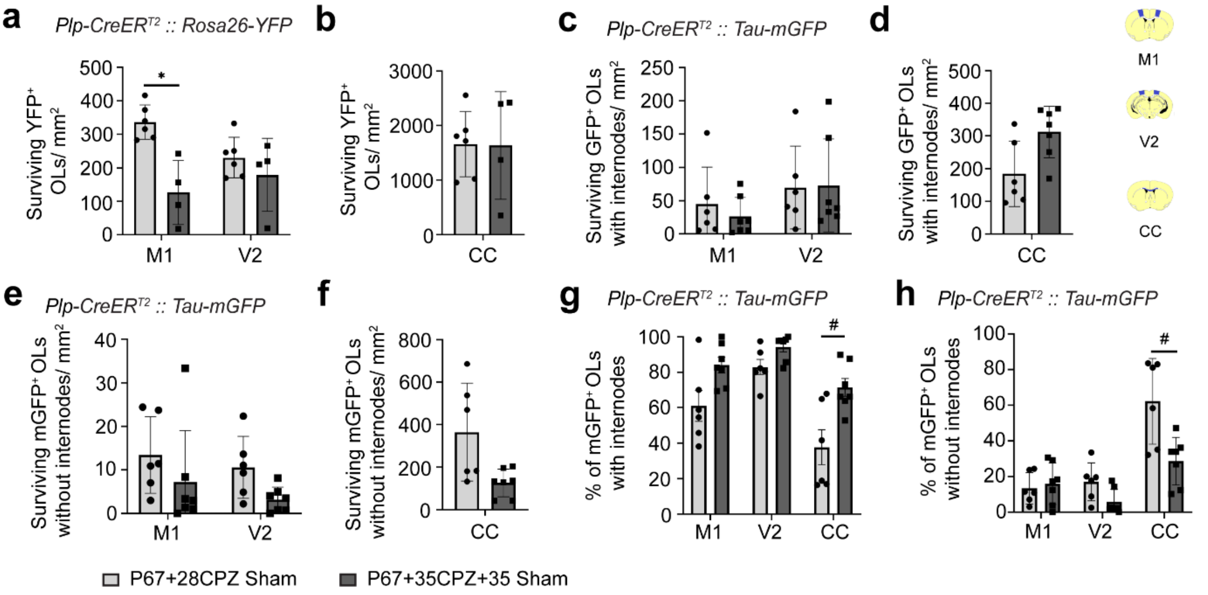


***Figure S4: OL loss occurs more slowly in M1 than V2 or the CC with CPZ feeding, but more of the surviving CC OLs transition from being completely demyelinated to supporting new internodes after CPZ withdrawal.***

**a-b.** The density of surviving YFP$^+$ OLs in M1, V2 (**a**) and the CC (**b**) of P67+28CPZ *Plp-CreER$^{T2}$ :: Rosa26-YFP* sham mice (n=6) or P67+35CPZ+35 *Plp-CreER$^{T2}$ :: Rosa26-YFP* sham mice (n=4). Repeated measures two-way ANOVA with Geisser-Greenhouse correction: treatment F (1, 8) = 0.24, p = 0.64; region F (1.0, 8.1) = 39.98, p = 0.0002; interaction F (2, 16) = 0.15, p = 0.86. **c-d.** The density of surviving mGFP$^+$ OLs with internodes in M1, V2 (**c**) and the CC (**d**) of P67+28CPZ *Plp-CreER$^{T2}$ :: Tau-mGFP* sham mice (n=6) and P67+35CPZ+35 *Plp-CreER$^{T2}$ :: Tau-mGFP* sham mice (n=7). Repeated measures two-way ANOVA with Geisser-Greenhouse correction: treatment F (1, 11) = 1.34, p = 0.271; region F (1.26, 13.86) = 83.6, p < 0.0001; interaction F (2, 22) = 10.14, p = 0.0008. **e-f.** The density of surviving mGFP$^+$ OLs with internodes in M1, V2 (**e**) and the CC (**f**) of P67+28CPZ *Plp-CreER$^{T2}$ :: Tau-mGFP* sham mice (n=6) and P67+35CPZ+35 *Plp-CreER$^{T2}$ :: Tau-mGFP* sham mice (n=7). Repeated measures two-way ANOVA with Geisser-Greenhouse correction: treatment F (1, 11) = 7.91, p = 0.017; region F (1.004, 11.05) = 27.11, p = 0.0003; interaction F (2, 22) = 6.48, p = 0.0061. **g.** The proportion (%) of surviving mGFP$^+$ OLs with internodes in M1, V2 and the CC of P67+28CPZ *Plp-CreER$^{T2}$ :: Tau-mGFP* sham mice (n=6) and P67+35CPZ+35 *Plp-CreER$^{T2}$ :: Tau-mGFP* sham mice (n=7). Repeated measures two-way ANOVA with Geisser-Greenhouse correction: treatment F (1, 11) = 10.92, p = 0.007; region F (1.788, 19.67) = 28.31, p < 0.0001; interaction F (2, 22) = 3.047, p = 0.068. **h.** The proportion (%) of surviving mGFP$^+$ OLs without internodes in M1, V2 and the CC of P67+28CPZ *Plp-CreER$^{T2}$ :: Tau-mGFP* sham mice and P67+35CPZ+35 *Plp-CreER$^{T2}$ :: Tau-mGFP* sham mice. Repeated measures two-way ANOVA with Geisser-Greenhouse correction: treatment F (1, 11) = 8.99, p = 0.012; region F (1.314, 14.46) = 27.24, p < 0.0001; interaction F (2, 22) = 6.421, p = 0.0064. Schematics of coronal brain sections show the analysed regions in blue i.e. M1 (~ Bregma +0.5), V2 (~ Bregma -2.5) and CC underlying M1 (~ Bregma +0.5). Data are presented as mean ± SD. Bonferroni post-test: *p < 0.05, # p = 0.052.
